# Non-coding exon splicing orchestrates tissue-specific expression of two functionally distinct mitochondrial isoforms of Mitofusin 2

**DOI:** 10.1101/2025.09.26.678759

**Authors:** Claudine David, Chloé Barsa, Camille Evain, Annika Krüger, Marion Papin, Rose Bernadet, Guillaume Duranthon, Thibaut Molinié, Elodie Cougouilles, Jim Dompierre, Manuel Rojo, Antoine Lefevre, Philippe Pasdois, Joanna Rorbach, Bianca H Habermann, Arnaud Mourier

**Affiliations:** University of Bordeaux, CNRS, IBGC, UMR 5095, F-33000 Bordeaux, France; Department of Medical Biochemistry and Biophysics, Division of Molecular Metabolism, Karolinska Institutet, Biomedicum, 171 65 Solna, Sweden; Max Planck Institute Biology of Ageing, Karolinska Institutet Laboratory, Karolinska Institutet, Stockholm, Sweden; Niche, Nutrition, Cancer & Oxidative metabolism (N2COx), UMR 1069, INSERM, University of Tours, Tours, France; Université de Tours, INSERM, Imaging Brain & Neuropsychiatry iBraiN U1253, 37032, Tours, France; Plateforme de Métabolomique et d’Analyses Chimiques, US61 ASB, Université de Tours, CHRU Tours, Inserm, Tours, France

**Keywords:** Mitochondria, Mitochondrial dynamics, Fusion, Mitofusin 2, Bioenergetics, Alternative ATG, Alternative translation initiation, 5’UTR alternative splicing

## Abstract

In mammals, outer mitochondrial membrane fusion controls mitochondrial network morphology, promotes subcellular trafficking and ensures proper mitochondrial content mixing. This process is controlled by two homologous dynamin related proteins, MFN1 and MFN2, which are ubiquitously expressed across tissues. Intriguingly, in humans, loss of function of MFN2 is responsible for the Charcot-Marie-Tooth type 2A neuropathy, whereas no pathogenic mutations are known in MFN1. Here, we report a short isoform of MFN2 that is generated by an evolutionary conserved alternative translation initiation site. This short mitochondrial MFN2 isoform (S-MFN2) is highly abundant in the mouse brain and its tissue-specific expression is controlled by the splicing of non-coding 5’exon(s), a mechanism conserved in humans. The S-MFN2 isoform has lower mitochondrial fusogenic capacity but seems critical for the trafficking and storage of neutral lipid. The genetically controlled tissue-specific ratio between L-MFN2 and S-MFN2 isoforms sheds important new light to understand both the tissue- and isoform-specificities of the MFN2 driven pathologies.

## INTRODUCTION

Mammalian mitochondria are dynamic organelles, present as a long interconnected tubular network or as individual, fragmented units that may undergo intracellular transport^1,2^. A family of dynamin-related GTPases regulate mitochondrial morphology through fission and fusion of the mitochondrial membranes^3–6^. The dynamin-related protein 1 (DRP1) mediates division of mitochondria, mitofusin 1 and 2 (MFN1 and MFN2) control outer mitochondrial membrane fusion (OMM), whereas optic atrophy 1 (OPA1) controls inner mitochondrial membrane (IMM) fusion. The physiological importance of mitochondrial fusion has been demonstrated by the fact that the above mentioned fusion proteins are all essential for mouse embryonic development^3,8^, and mutations in *MFN2* or *OPA1* respectively cause neuropathy^10–12^ and optic atrophy^13,14^ in humans. The pathological spectrum associated with disturbed mitochondrial dynamics has recently expanded to include Parkinson’s, Huntington’s and Alzheimer’s diseases^15–17^. Despite the high similarity (81%) between the paralogous proteins MFN1 and MFN2, and their common role in mitochondrial fusion through their physical interaction, CMT2A neuropathies are exclusively associated with mutations in *MFN2*. The reason why CMT2A is only caused by mutations in *MFN2* remains elusive and puzzling given the structural and functional similarities between MFN1 and MFN2.

Due to their key role in energy transduction, mitochondria localize at sites of high energy demand and mitochondrial fusion is therefore finely adjusted to the shape and bioenergetic needs of different cell types. Tissue-specific adjustment of mitochondrial fusion is also key to coordinating mitochondrial activity with tissue-specific needs for mtDNA replication and maintenance^18,19^. However, the mechanisms adjusting OMM fusion to cell type-specific needs are not understood. In contrast, OPA1, which controls IMM fusion, is expressed as at least 8 alternate mRNA splice isoforms, and in addition, the OPA1 protein undergoes processing with different proteases to regulate its activity in a tissue-specific manner^13,20^. Recently, long-range splicing of mRNAs encoding *MFN2* has been reported^21^ leading to the production of small MFN2 protein isoforms lacking the GTPase domain. These short MFN2 protein isoforms are present at very low levels, and targeted to the endoplasmic reticulum and have no role in mitochondrial fusion. Here, we demonstrate that a highly conserved alternative translation initiation site produces a MFN2 isoform presenting a 20 AA shorter amino terminus (S-MFN2) compared to the larger MFN2 form (L-MFN2). Intriguingly, despite having been previously observed in a few studies, these two closely running MFN2 protein forms remained unaddressed. Interestingly, the tissue-specific expression of the L-MFN2 and S-MFN2 isoforms is orchestrated through alternate splicing of 5’non-coding exons during differentiation, notably correlated with abundance of the slicing factor RBM24. The S-MFN2 isoform is at least five times more abundant than L-MFN2 in many tissues, reaching more than 90% of the total mitochondrial MFN protein levels in brain. Both L- and S-MFN2 forms locate as foci at the tips and branches of the mitochondrial OM, and expose their C and N terminus to the cytoplasm. Multimodal functional analyses, characterizing mitochondrial dynamics, genetics and bioenergetics, demonstrated that S-MFN2 displays reduced fusogenic activity compared to the L-MFN2. Interestingly, lipidomic analyses unravelled that mitochondrial levels of neutral lipids derived from the mevalonate pathway and lipid droplets are specifically modulated in response to S-MFN2 expression. The novel genetic regulation controlling the ratio between L-MFN2 and S-MFN2 across tissues provides an original mechanism for tissue-specific control of mitochondrial dynamics.

## RESULTS

### Non-coding exon splicing controls expression of MFN2 isoforms

The development of high-resolution denaturing electrophoresis and western blot analyses for MFN2, which combines long electrophoretic migration and fluorescent immunolabelling, unravelled the presence of a second MFN2 band, which was undetectable under classical western blot conditions (Fig. 1A). The use of independent anti-MFN2 antibodies targeting different epitopes (Supplementary Fig. S1A), together with tissue-specific *Mfn2* heart and skeletal muscle conditional KO^22,23^ (Fig. 1A), unambiguously demonstrated that these two protein entities resolved by western blot are encoded by *Mfn2* gene. Interestingly, the high-resolution western blot analyses applied to total extract (Fig. 1B) and purified mitochondria (Supplementary Fig. S1B) of different mouse tissues, showed that the respective abundancy of the two MFN2 entities strongly differs across tissues. In contrast to heart and skeletal muscle, where both isoforms seem equally present, liver, kidney and brain mainly express the low molecular weight MFN2 entity and the high molecular weight is hardly detected. To investigate the mechanism generating these MFN2 entities presenting close apparent molecular weight, we first decided to design various sets of primers to detect potential short-range splicing within *Mfn2* transcripts. To this end, we targeted cDNA generated from tissues or mouse embryonic fibroblasts (MEFs) with a combination of primers, designed to be evenly distributed along *Mfn2* mRNA. None of the primer combinations could identify short-range splicing in the protein coding region (exons 4-20) (Fig. 1C; Supplementary Fig. S1D). Intriguingly, the PCR-based detection of the splicing variant, combined with sequencing analyses, unambiguously demonstrated that the non-coding Exon 3 localized in *Mfn2* 5’UTR was present in heart but spliced in mouse embryonic fibroblasts (MEFs) (Fig. 1D). Extending this analysis to different tissues demonstrated that Exon 3 was present both in heart and skeletal muscle but was almost undetectable in liver, kidney and brain (Fig. 1E). The discovery of this *Mfn2* tissue specific 5’exon splicing prompted us to validate this observation in existing bulk RNA sequencing data of different mouse tissues^24^. Bioinformatic analyses validated the occurrence of exon 3 splicing of the 5’ UTR in brain, liver, kidney and various adipose tissues, and retention of this exon in heart, skeletal muscle and lung (Supplementary Fig. S1 E, F).

**Figure 1.**
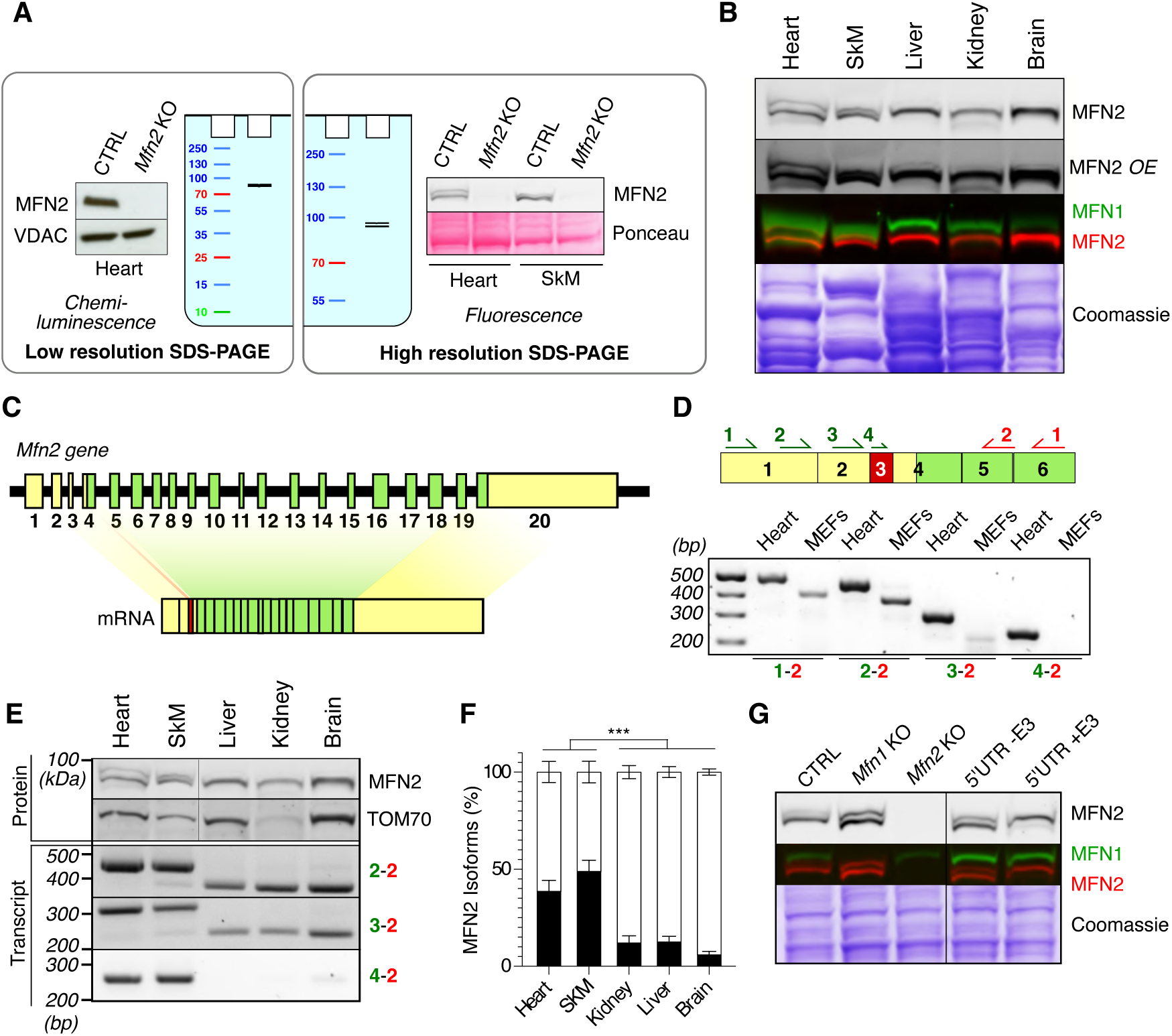
Tissue-specific MFN2 doublet profiles correlate with the alternative splicing of the non-coding Exon 3. (A) Illustration comparing the low- and high-resolution denaturing electrophoreses and western blot protocols, and their respective capacity to detect MFN2 doublets in total protein extracts form mouse heart and skeletal muscle. (B) MFN2 proteins doublet detection by high-resolution denaturing electrophoresis and western blot performed with total protein extracts from the indicated mouse tissues (n ≥ 7 per tissue, *OE* abbreviation means Over-Exposed). (C) Schematic representation of *Mfn2* mouse gene and transcript. Non coding exons (5’UTR and 3’UTR) appear as yellow boxes, and coding exons as green boxes. (D) PCR detection of *Mfn2* splice variants performed on cDNA purified from mouse Heart and MEFs. Forward primers (green half arrows and numbers) and reverse primers (red half arrows and numbers) are positioned on *Mfn2* cDNA. Non coding and coding regions respectively appear in yellow and green, whereas the spliced Exon 3 appears as a red box. The combination of primers used for the reaction is indicated underneath each PCR product (n ≥ 3 for each PCR combinations) (E) High-resolution denaturing electrophoresis and western blot of MFN2 and TOM70, performed on total protein extracts from indicated tissues (upper part) and *Mfn2* transcript splice variants (lower part) characterized from cDNA of the indicated mouse tissues (n ≥ 7 mouse tissues extracts and n ≥ 3 cDNAs). (F) Proportion of upper (black bars) and lower (white bars) proteins present in the MFN2 doublet, determined by densitometric analyses of high-resolution western blot analyses performed on total protein extracts from the indicated tissues (n ≥ 7 mouse tissues). Error bars ± SEM (***p<0.0001). (G) MFN2 protein doublet detection by high resolution denaturing electrophoresis and western blot in *Mfn2* KO stable MEF lines expressing *Mfn2* ORF preceded by 5’UTR containing (+E3) or not (- E3) Exon 3 (n ≥ 6 MEFs independent experiments).

The occurrence of the exon 3 alternative splicing strikingly correlated with the change in abundancy between the two MFN2 entities. The ratio between both MFN2 protein entities shifts from a situation, in heart and skeletal muscles where both entities are co-expressed when exon 3 is present, to a severe decrease of the high molecular weight MFN2 entity level in tissues where exon 3 is spliced (Fig. 1E, F). This observation prompted to generate new transgenic cell lines to specifically investigate the role of the 5’exon 3 on the control of MFN2 isoforms levels. To this end, we transduced *Mfn2* KO MEFs with retrovirus encoding the *Mfn2* open reading frame preceded by its 5’UTR containing (5’UTR+E3 *Mfn2*), or not, exon 3 (5’UTR-E3 *Mfn2*). This approach was validated by the fact that MFN2 entities levels (high versus low molecular weight) observed in stable *Mfn2* KO cells transduced with 5’UTR-E3 *Mfn2*, faithfully recapitulate the levels observed in control MEFs (where exon 3 is spliced) (Fig. 1G). Remarkably, alike heart and skeletal muscle, the presence of exon 3 in *Mfn2* ORF (5’UTR+E3 *Mfn2*), strongly promoted the expression of the high molecular weight MFN2 entity in transduced MEFs (Fig. 1G). Altogether, our data demonstrate that alternative splicing of the non-coding exon 3 localized in the 5’UTR region of *Mfn2* actively regulates the proportion between two MFN2 protein entities.

### The short MFN2 isoform originates from a second functional alternative translation initiation site conserved in mouse and human

Taking a close look at the *Mfn2* 5’UTR sequence, we noticed the presence of a second ATG localized 60bp downstream the canonical ATG of *Mfn2*. As presented in Figure 2 A and C, when we performed a protein pairwise sequence alignment between mouse and human MFN1 and MFN2, we noticed that a second methionine located 20AA downstream of the starting methionine of MFN2 was aligned with the starting methionine of MFN1. The position of this second methionine is highly conserved in vertebrates ranging from xenopus, zebrafish to mammals (Fig. 2A, C; Supplementary Fig. S2A). The concordance between the small electrophoretic shift separating MFN2 isoforms in Western blot (Fig. 1B) and the shift in molecular weight provoked by the loss of 20AA (∼2.4 kDa) hypothetically observed if the Met21 could be used as a start codon, prompted us to evaluate the functionality of this second ATG site. To investigate the role of Met1 and Met21 on MFN2 isoforms expression, we generated stable *Mfn2* KO MEFs transduced with *Mfn2* ORF subjected to targeted mutagenesis (ATG-Met mutated to ATC-Ile). Interestingly, high-resolution electrophoresis performed with total protein extracts demonstrated that upper and lower MFN2 entities were respectively lost in Met1 knock-in MEFs (*Mfn2* ΔATG1), and Met21 knock-in MEFs (*Mfn2* ΔATG2) (Fig. 2B). This experiment demonstrated that the highly conserved second ATG site is active, and that this downstream translation initiation site is generating a 20AA shorter MFN2 isoform. We named these long and short products L-MFN2 and S-MFN2.

**Figure 2.**
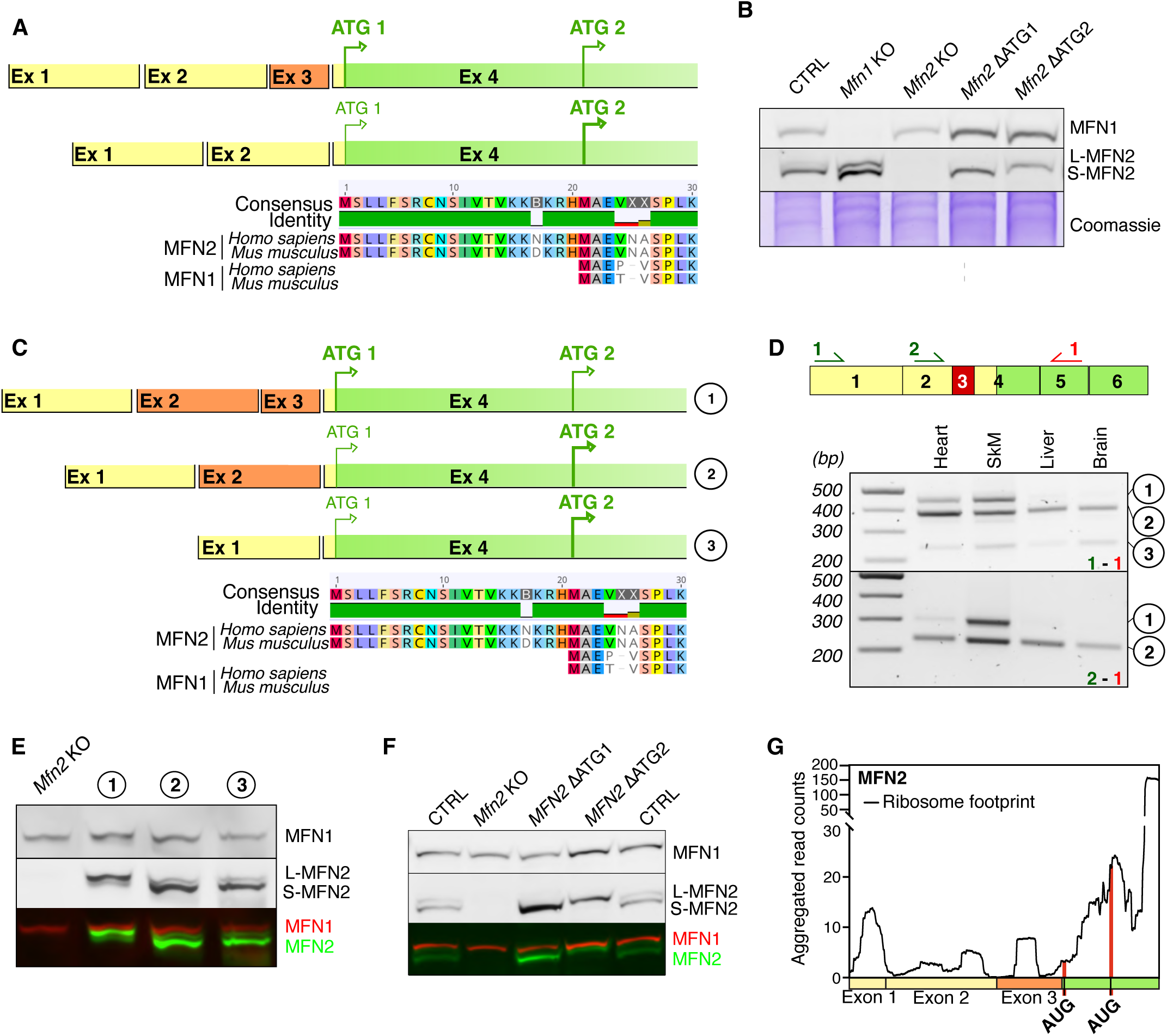
Human and mouse *MFN2* 5’UTR exon(s) splicing orchestrate(s) MFN2 alternative translation initiations in mouse and human. (A) (top) Schematic representation of mouse *Mfn2* 5’UTR presenting non-coding (yellow) and coding region (green) and exon undergoing tissue specific alternatively splicing (orange). (bottom) The multiple protein sequence alignments performed between human and mouse MFN1, and MFN2 unravel the presence of a conserved and potentially active alternative translation initiation site (ATG2). (B) High-resolution denaturing electrophoresis and western blot of MFN2 performed with total protein extracts from control, *Mfn1* and *Mfn2* KO and stable MEF lines transduced with *Mfn2* ORF presenting disrupted ATG1 or ATG2 by targeted mutagenesis (representative n ≥ 3 independent experiments). (C) (top) Schematic representation of human *Mfn2* 5’UTR presenting non-coding (yellow) and coding region (green) and exons undergoing tissue specific alternatively splicing (orange). (bottom) The multiple protein sequence alignments performed between human and mouse MFN1, MFN2 unravel the presence of a conserved and potentially active alternative translation initiation site (ATG2). (D) PCR detection of *MFN2* splice variants performed on cDNA of the indicated human tissues. Forward (green) and reverse (red) primers used are positioned on *MFN2* cDNA. The three different splice variants (1, 2 and 3) validated by sequencing and defined in (C) are labelled on the right side (n ≥ 3). (E) Detection by high-resolution denaturing electrophoresis and western blot of human MFN2 in total protein extracts from stable *Mfn2* KO MEF lines transduced with *MFN2* ORF preceded by 5’UTR variants defined in C. (n ≥ 6 MEF total protein extracts). (F) High-resolution denaturing electrophoresis and western blot of MFN2 performed with total protein extracts from *Mfn2* KO and stable MEF lines transduced with the human *MFN2* ORF presenting disrupted ATG1 or ATG2 by targeted mutagenesis (representative n ≥ 3 independent experiments). (G) Alignment of existing ribo-seq data from mouse heart tissue to MFN2 transcripts ^88^. Region 1-400 nucleotides is shown. Coding, non-coding region, and exon 3 alternatively spliced are coloured in green, yellow and orange respectively. Locations of translation initiation codons ATG are presented with a red line.

To investigate if the Met21 was also active and able to generate S-MFN2 in humans, we subjected the total extract of Hela cells and *MFN2* KD to high-resolution electrophoresis and could observe that in contrast to MFN1, MFN2 appeared as a double band, and that both L-MFN2 and S-MFN2 consistently disappeared in *MFN2* KD cells (Supplementary Fig. S2C). This observation prompted us to characterize the human *MFN2* 5’UTR using a similar strategy as the one developed in mouse. To identify short-range spliced MFN2 variants, cDNA purified from human tissues were subjected to PCR-based detection of splice variants, combined with sequencing analyses. In line with the *Mfn2* transcript in mouse, our analyses did not identify exon splicing within the coding region of the human *MFN2* transcript (Supplementary Fig. S2B). We observed that human and mouse *Mfn2* 5’UTR are very similar, containing 3 exons, and that the non-coding 5’exon 3 can undergo alternative splicing. However, in contrast to the mouse *Mfn2*, the human exon 3 of the 5’ UTR can be spliced alone or in combination with 5’exon 2 (Fig. 2C, D). The tissue-specific pattern of alternative splicing is remarkably conserved, as the *MFN2* transcript variant possessing the 5’exon 3 is found preferentially expressed in heart and skeletal muscle whereas liver and brain almost exclusively expressed spliced variants (Fig. 2D and E). To further investigate the impact of these different 5’UTRs on the expression of MFN2 isoforms, we transduced *Mfn2* KO cells using a vector containing the human *MFN2* ORF preceded by the three different 5’UTR splicing variants. High-resolution electrophoresis analyses of total cell extract clearly demonstrated that the loss of the non-coding 5’ exon 3 alone or associated with 5’ exon 2 downregulates the expression of the L-MFN2, promoting the S-MFN2 expression levels (Fig. 2E). The alternative ATG was also confirmed in human ORFs by generating stable cell lines expressing human *MFN2* ORFs with mutated ATG1 or ATG2 (Fig. 2F). High resolution, denaturing electrophoresis and western blot clearly confirmed that human *MFN2* ATG2 is an active translation initiation site producing the S-MFN2 isoform. These results demonstrate that splicing of non-coding 5’UTR exon(s), modulating the expression of L and S-MFN2, is conserved between mouse and human.

To learn more about the translational regulation of MFN2 expression, we reanalysed existing ribo-seq footprinting data^25^. The alignment of footprints on MFN1 transcript confirmed the usage of the canonical AUG start site on this transcript in mouse heart tissue (Supplementary Fig. S2D). Furthermore, the low number of reads in the 5’-UTR region suggests that this region is not actively translated. In contrast, the alignments of footprints on MFN2 transcript presenting 5’exon 3 unravelled an active interplay between *Mfn2* 5’UTR exons and ribosomes (Fig. 2G). We not only observed a significant accumulation of reads in exon 1, exon 2, and exon 3 regions, which supports our previous analyses of transcripts showing the presence of 5’exon 3 in heart (Fig. 1D, E), but we also observed a predominant accumulation of reads on the canonical as well as the second AUG, which supports our conclusion that the second AUG start codon is functional (Fig. 2G). To further investigate the role of the 5’UTR in the regulation of MFN2 isoform expression and its conservation in human, we specifically looked at translation initiation sites in HEK293T cells on MFN1 and MFN2 transcripts via a recently developed approach (TISCA = translation initiation site detection by translation Complex Analysis)^26^. The approach includes a combinatorial analysis of ribosome footprints from 40S subunits (=40S-seq) and ribosome footprints from lactimidomycin (LTM) treated cells (= GTI-seq (global translation initiation)). Start sites are indicated by a combined presence of a GTI-seq peak and a decrease of 40S reads. Alignment of both datasets to MFN1 transcript confirmed the usage of the canonical AUG start codon (Supplementary Fig. S2E). Even though the generally low coverage of reads (in particular from GTI-seq data) does not allow detailed conclusions, the alignment of GTI-seq and 40S-seq data to MFN2 supported our previous findings showing that translation initiation signals at ATG2 are stronger than ATG1 in the absence of exon 3 (no read from GTI-seq data covering ATG1) (Supplementary Fig. S2F). Several regions of active translation were detected in non-coding 5’exons that colocalise to the upstream ORF (uORF) sequences identified by uORFdb (green bands Supplementary Fig. S2E, F) ^27^. The uORFs have received increasing interest as it has been recently demonstrated that they act as regulators repressing or even supressing the expression of the downstream ORF ^28–30^. The high number and activity of uORFs present in human and mouse MFN2 (Fig. 2G; Supplementary Fig. S2F) could, in synergy with 5’exon(s) splicing, contribute to the control of the expression of both MFN2 isoforms.

To investigate if the splicing and therefore the MFN2 isoforms expression could be regulated by metabolic stresses, control MEFs were subjected to various metabolic conditions known to challenge mitochondrial dynamics^23,31,32^. The analyses showed that cultivating cells under different carbon sources or serum deprivation during three days, did not induce significant changes in the expression of MFN2 isoforms and did not alter the morphology of mitochondrial network (Supplementary Fig. S3A-C). To investigate the interplay between MFN2 isoforms and metabolism, we assessed S- and L-MFN2 expression in various muscles, including the heart, quadriceps, diaphragm, soleus (SOL), and extensor digitorum longus (EDL), which have distinct energy metabolism profiles *i.e.* soleus being slow/oxidative and EDL being fast/glycolytic^33^ (Supplementary Fig. S3D, E). Denaturing electrophoresis and Western blot analysis revealed similar S- and L-MFN2 expression levels across muscle types (Supplementary Fig. S3D, E). Altogether, our investigation demonstrated that the MFN2 mitochondrial isoforms expression does not respond to specific metabolic conditions but seems to be orchestrated during cells and tissues differentiations. In line with this, a work from Jing Liu and colleagues has recently characterized a mouse cardiac conditional knockout of the RNA binding motif protein 24 (RBM24), a key component of the splicing machinery involved in cardiac muscle differentiation and convincingly unraveled that the loss of RBM24 was increasing the level of cardiac Mfn2 transcripts spliced for the exon 3^34^. Consistent with this finding, we performed denaturing gel electrophoresis and western blot analyses (Supplementary Fig. S3D-F) and, despite faint traces detected in total brain samples (Supplementary Fig. S3F), our analyses demonstrated that RBM24 is not exclusively expressed in heart but is also highly expressed in various muscle tissues known to present a 5’UTR with exon 3 (Fig. 1D, E; Fig. 2D) and co-expressing both the S- and L-MFN2 isoforms (Fig. 1A, B, F; Supplementary Fig. S3D-F). Our findings support the study by Jing Liu *et al.* ^34^ and suggest that RBM24’s role in preventing the splicing of exon 3 of *Mfn2* mRNA may extend beyond cardiac tissue to include other muscle types.

### S-MFN2 is a mitochondrial isoform very highly expressed in brain

Shortening of the proteins N-terminal domain, secondary to alternative translation initiation, was proposed to constitute a mechanism modulating the subcellular targeting of proteins isoforms^35,36^. To characterize the subcellular localisation of S-MFN2, we engineered *Mfn2* KO MEFs expressing L-MFN2 and S-MFN2 carrying a C-terminal fluorescent tag (EGFP or mCherry), a strategy known not to alter the mitochondrial localisation of MFN2^37,38^. Epifluorescence microscopy experiments demonstrated that S-MFN2 and L-MFN2 isoforms clearly colocalise. Their mitochondrial localisation was validated using the well-established VDAC staining (Fig. 3A). To characterize with greater detail the mitochondrial distribution of MFN2 isoforms, we adapted Ultrastructure Expansion Microscopy (U-ExM) to mitochondrial analyses allowing us to improve the resolution down to 40 nm^39^. The U-ExM imaging approach unravelled that the OMM distribution patterns between the S-MFN2 and VDAC were clearly distinct. In contrast to VDAC, S-MFN2 appeared often enriched at the tips of mitochondria or at sites of inter mitochondrial contact (Fig. 3B). U-ExM analyses performed with cells co-expressing tagged MFN2 isoforms demonstrated that S-MFN2 and L-MFN2 distributions are distinct with only partial overlap in the periphery of tubular areas (Fig. 3C). However, both MFN2 isoforms accumulated focally at mitochondrial tips and branches, areas commonly defined as fusion and fission ‘hot spot’ sites^40,41^. We then decided to independently confirm the mitochondrial targeting of MFN2 isoforms in heart tissue, where endogenous S-MFN2 and L-MFN2 are evenly expressed (Fig. 1B). To this end, we performed subcellular fractionation using differential centrifugations combined with percoll^®^ density gradients, allowing us to obtain independent fractions presenting various degrees of mitochondrial enrichment. The western blot analyses of these different fractions confirmed that the S-MFN2 fractionation pattern was identical to L-MFN2 and MFN1. In line with microscopy analyses performed in MEFs (Fig. 3A-C), the majority of MFN2 isoforms were found predominantly enriched with mitochondrial markers (TFAM, TOM22) (Fig. 3D). Then, we investigated if the shortening of N-terminal extension in S-MFN2 isoforms could impact its topology. To this end, we subjected isolated mitochondria purified from heart or MEFs, to classical protease shaving protocols followed by Western blotting to evaluate protease accessibility of MFN2 Nter and Cter extremities (Fig. 3E).

**Figure 3.**
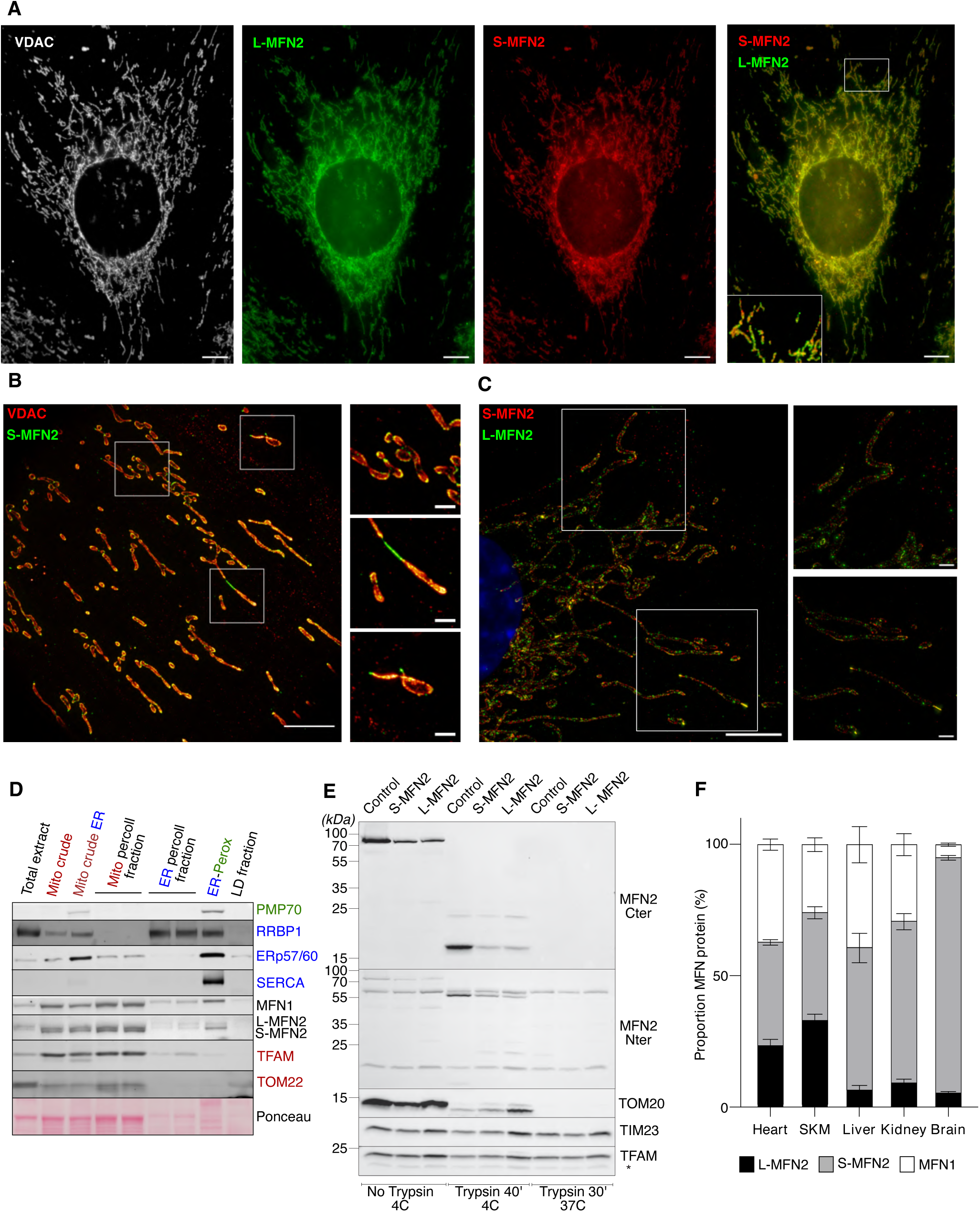
S-MFN2 is the most abundant mitochondrial isoform and exhibit a canonical MFN2 topology. (A) Representative epifluorescence microscopy images of *Mfn2* KO MEFs expressing L-MFN2 (green) and S-MFN2 (red) tagged with fluorescent proteins colocalized with the mitochondrial protein through immunostaining analyses performed using anti-VDAC antibody (grey). (scale bar = 10µm) (B) Representative expansion microscopy of *Mfn2* KO MEFs S-MFN2 tagged with fluorescent proteins (green) colocalized with the mitochondrial protein through immunostaining analyses performed using anti-VDAC antibody (red). (scale bar = 10µm and 2µm in insert) (C) Representative expansion microscopy of *Mfn2* KO MEFs co-expressing tagged L-MFN2 (green) and S-MFN2 (red) tagged with fluorescent proteins. (scale bar = 10µm and 2µm in insert) (D) Mouse heart subcellular fractionation following MFN1, MFN2 isoforms, ER, mitochondria and peroxisomes protein markers. The overall enrichment of MFN isoforms in the different fractions: Mito crude (pellet 1 000G), Mito crude ER (pellet 7 000G), ER-Perox (Pellet at 30 000G) and Low density -LD (supernatant at 30 000G) as well as Mito percoll^®^ and ER percoll^®^ (respectively lower and upper phase in 20% percoll^®^ at 10 000G performed on Mitocrude ER extract), were analysed by high resolution electrophoresis and western blot. (representative n ≥ 3 independent experiments) (E) MFN2 isoforms topology determined using classical trypsin protease shaving assay performed on isolated mitochondria from control and *Mfn2* KO MEFs expressing L-MFN2 and S-MFN2 (representative of n = 4 independent experiments, * indicates aspecificity). (F) Densitometric quantification of the MFN1 (white), L-MFN2 (black) and S-MFN2 (grey) proportion across the indicated mouse tissues total protein extracts. Quantification was achieved thanks to MFN1 and MFN2 standards produced in E.coli. (n ≥ 5 independent experiments).

Sequential solubilisation of the OMM and IMM membrane using increasing digitonin to protein ratio demonstrated that TOM20 (extra mitochondrial – OMM anchored), TIM23 for (intermembrane space - IMM anchored) and TFAM (intramitochondrial - matrixial) were reliable markers to evaluate the submitochondrial localisation and topology of proteins (Supplementary Fig. S4A). The analysis presented in Figure 3E demonstrated that the MFN2 Cter and Nter, partially digested at 4°C, were in fact totally digested by trypsin when the protease shaving assay was performed at 37°C and in the absence of detergent. The integrity of the OMM at 37°C was validated by the intactness of TIM23 (IMS) and TFAM (matrix) markers, even after 30 minutes incubation (Supplementary Fig. S4B, C). This experiment performed with higher time resolution in control MEFs (Supplementary Fig. S4B) and heart mitochondria (Supplementary Fig. S4C) demonstrated that MFN2 N and C termini, of both MFN2 isoforms behave like TOM20 and consequently are exposed toward the cytosolic face of the OMM. To tackle the rationale of having different mitofusin mitochondrial isoforms, we then developed a MFN-targeted quantitative western blot assay using MFN1 and MFN2 protein standards produced in *E.coli*, to quantify the tissue-specific distribution of the three mitochondrial MFN proteins. In line with the results obtained with total extracts of tissues (Fig. 1E, F), the quantification of MFN1, L-MFN2 and S-MFN2 proportion performed on total extracts (Fig. 3F) or isolated mitochondria (Supplementary Fig. S4D, E) unravelled that S-MFN2 is a very abundant isoform. In fact, S-MFN2 is the most highly expressed MFN proteins in heart and skeletal muscle (∼40%), liver (∼55%), kidney (∼60%) and, remarkably, represent 90% of the MFN proteins expressed in brain. The extremely low levels of MFN1 and L-MFN2 compared to S-MFN2 observed in brain was highly significant when compared with other tissues. These analyses demonstrate that S-MFN2 is a very abundant mitochondrial mitofusin anchored in the OMM, according to the canonical MFN topology.

### S-MFN2 is fusion-competent but presents a low efficiency compared to L-MFN2

The diverging tissue expression patterns between MFN2 isoforms prompted us to characterise their functional properties. We first generated *Mfn1 KO and Mfn2* KO MEFs using a strategy allowing to generate control (*Mfn1*^Lox/Lox^ or *Mfn2^L^*^ox/Lox^) and KO MEFs from the same embryo. In line with *Mfn1* or *Mfn2* KO MEF models previously generated^7,42^, we observed that the loss of MFN1 or MFN2 impaired the mitochondrial network morphology (Supplementary Fig. S5A, B), but was not significantly affecting the morphology and distribution of other organelles as peroxisome or endoplasmic reticulum (Supplementary Fig. S5A; Supplementary Fig. S7A-D). To functionally characterize and compare L- and S-MFN2, we decided to transduce *Mfn2* KO MEFs with retrovirus expressing murine ORFs L-MFN2 (ΔATG2), S-MFN2 (ΔATG1). The different MEFs lines (Mfn2 KO, S-MFN2, L-MFN2) enabled the development of a single-cell analysis that directly correlates mitochondrial fusion activity to the expression levels of MFN2 isoforms. This approach, leveraging the heterogeneity of transgene expression inherent to virally transduced cells was designed to quantitatively assess mitochondrial network morphology (Fig. 4A-E) and mitochondrial DNA nucleoids content (Fig. 4F, G) per cell in regard of the expression level of each MFN2 isoforms. These analyses indicate that the morphology and mtDNA content were restored proportionally to the expression levels of each isoform (Fig. 4A-G). However, morphological parameters assessed through the ‘Mitochondria-Analyzer’ multiparametric analyses (Fig. 4B-E), together with the mtDNA nucleoids content (Fig. 4F-G), clearly demonstrated that the mitochondrial fusogenic activity of S-MFN2 is significantly lower than the L-MFN2 isoform. To investigate if this functional discrepancy between MFN2 isoforms was also present in human MFN2 isoforms we generated *Mfn1, Mfn2* dKO MEFs stably expressing human L- and S-MFN2^19,42,43^. Single-cell analyses conducted in independent fusion-deficient MEF models clearly demonstrated that human S-MFN2, like its murine counterpart, exhibited lower fusogenic activity compared to L-MFN2 (Supplementary Fig. S6A-G). To strengthen the conclusion obtained with ‘single cell’ analyses we also generated stable *Mfn* dKO MEFs cell lines expressing murine ORFs L-MFN2 (ΔATG2), S-MFN2 (ΔATG1) or both MFN2 isoforms (UTR-MFN2) preceded by the endogenous MEF 5’UTR (lacking exon 3) (Fig.5 ; Supplementary Fig. S5C). In line with the ‘single cell’ analyses performed on *Mfn* dKO MEFs expressing human MFN2 isoforms, the western-blot analyses performed on *Mfn* dKO MEFs expressing only one of the two MFN2 isoforms or coexpressing both isoforms (UTR-MFN2) demonstrated that the levels of S-Mfn2 is more than two times higher than the levels of L-MFN2 expressed alone or together with S-MFN2 (UTR-MFN2) (Fig. 5A, B). This very high expression of S-MFN2 assessed in *Mfn* dKO was consistent with the very high S-MFN2 expression level assessed in brain mitochondria, where the expression levels of both L-MFN2 and MFN1 are extremely low (Supplementary Fig. S5D). Despite the very high level of expression of S-MFN2, the important rescue of mitochondrial network morphology quantified in S-MFN2 dKO and L-MFN2 dKO were almost identical (Fig. 5C, D; Supplementary Fig. S5C). In line with this observation and despite their distinct expression levels, MFN2 isoforms were both able to completely restore mtDNA levels (Fig. 5E), and to counteract mtDNA clustering present in *Mfn* dKO (Fig. 5D; Supplementary Fig. S5C), and were partially restoring mitochondrial endogenous and complex I driven respiration in dKO MEFs (Fig. 5F). The mitochondrial network morphology and cell respiration were more efficiently restored in the 5’UTR *Mfn* dKO MEFs expressing both MFN2 isoforms, compared to *Mfn* dKO MEFs expressing only one of the two MFN2 isoforms, likely indicating a cooperative effect between L-MFN2 and S-MFN2 (Fig. 5C, D, F). The rescue of mitochondria and cell energy metabolism was also evidenced by the significant reduction of cell doubling time under High glucose or under conditions stressing mitochondrial activity as low serum or galactose (Fig. 5G). In line with the results obtain on mtDNA or mitochondrial bioenergetic rescue, the rescue of cell growth was always higher in line expressing L-MFN2 compared to S-MFN2. Altogether, our single cell and multimodal functional analyses unravelled that both L-MFN2 and S-MFN2 can restore mitochondrial morphology and activity through homotypic or heterotypic interactions. Interestingly, in regard to their respective expression levels, our analyses consistently demonstrated that the fusogenic activity of S-MFN2 is significantly lower than the L-MFN2.

**Figure 4.**
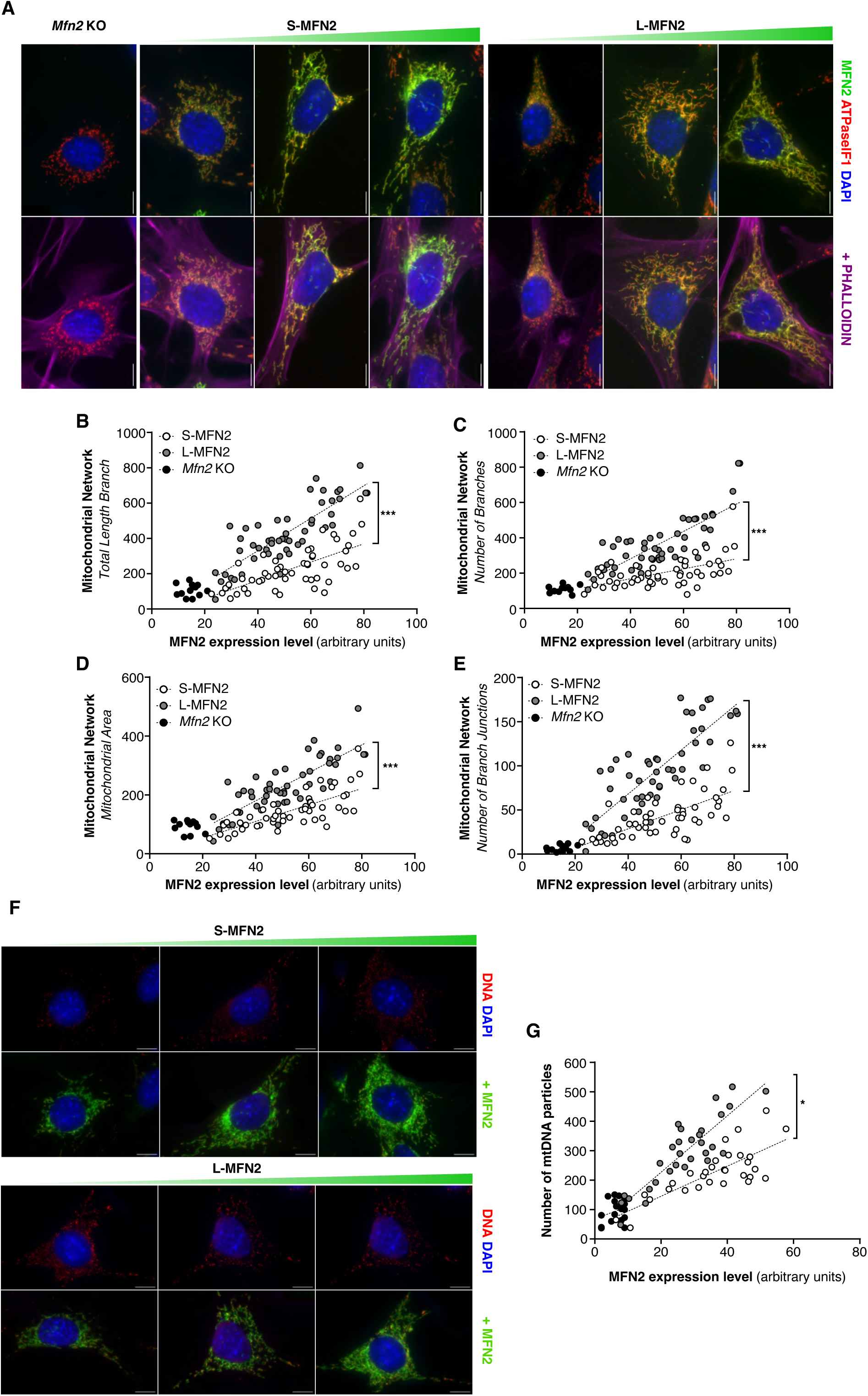
S-MFN2 has lower capacity to restore mitochondrial morphology and mitochondrial DNA levels than L-MFN2. (A) Representative epifluorescence microscopy of *Mfn2* KO MEFs expressing mouse S-MFN2 or L-MFN2, using anti-ATPase IF1 as a mitochondrial marker (red), anti-MFN2 D2D10 (green), and phalloidin as a cytoskeleton marker (magenta) (scale bar = 10 µm). (B-E) Mitochondrial morphology and network parameters (mitochondrial area, total length branch, number of branches and number of branch junctions) quantified with the MitoAnalyzer plugin and plotted against the MFN2 intensity on a single-cell basis. (*n* > 30 for *Mfn2* KO, *Mfn2* KO + S-MFN2 and *Mfn2* KO + L-MFN2 MEFs) PERMANOVA, the permutational multivariate analysis of variance test relative to *Mfn2* KO MEFs; *, P < 0.05; **, P < 0.01; ***, P < 0.001. (F) Representative epifluorescence microscopy of *Mfn2* KO MEFs expressing S-MFN2 or L-MFN2, using anti-DNA (red) to stain mtDNA particles and anti-MFN2 D2D10 (green) (scale bar = 10 µm). (G) The number of mtDNA particles per cell is plotted against MFN2 intensity for each analyzed cell. (*n* > 30 for *Mfn2* KO, *Mfn2* KO + S-MFN2 and *Mfn2* KO + L-MFN2 MEFs respectively) PERMANOVA, the permutational multivariate analysis of variance test relative to *Mfn2* KO MEFs; *, P < 0.05; **, P < 0.01; ***, P < 0.001.

**Figure 5.**
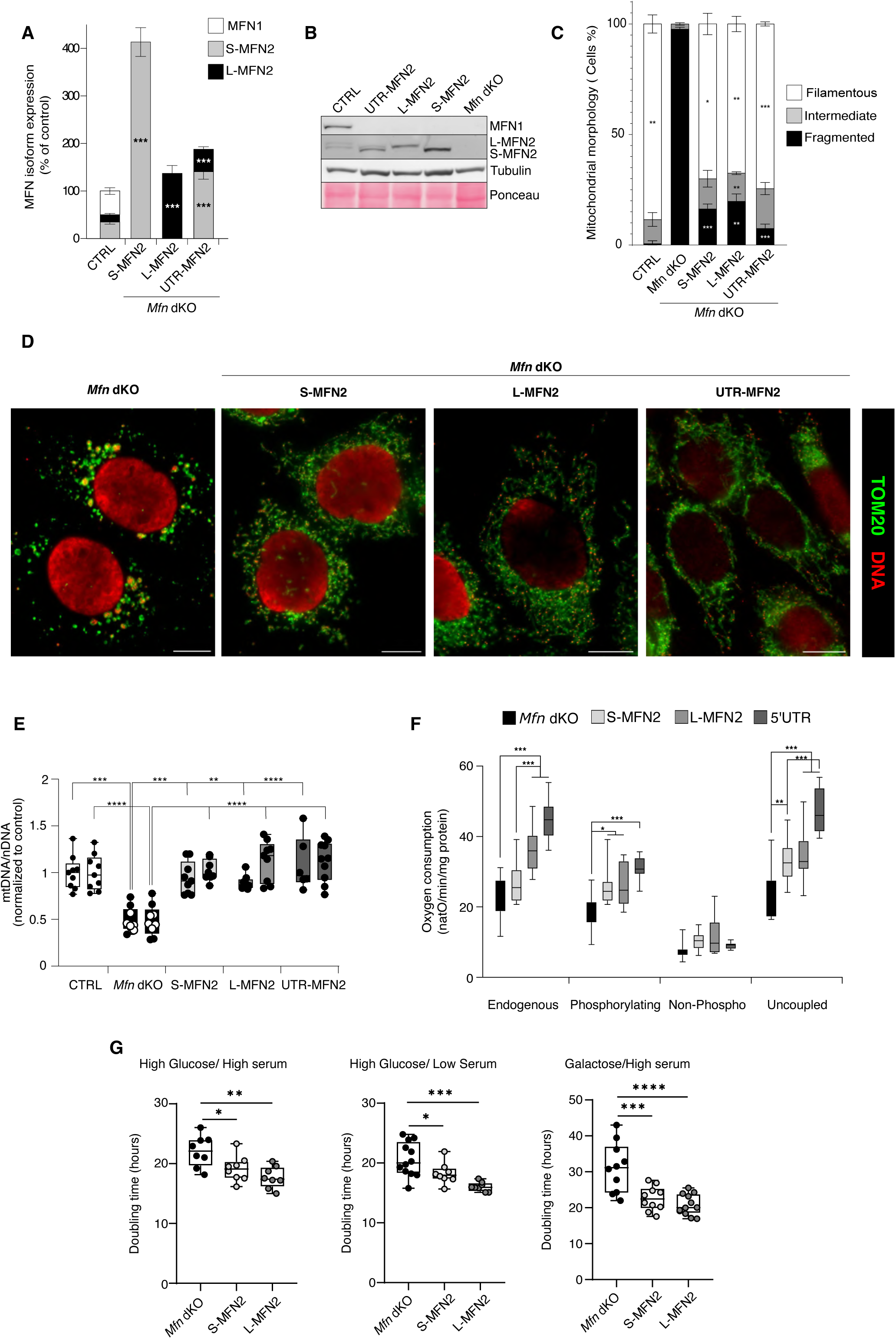
L-MFN2 and S-MFN2 isoforms present different expression profile and distinct capacities to restore mitochondrial morphology and bioenergetics. (A) Densitometric quantification of the MFN1 (white), L-MFN2 (black) and S-MFN2 (grey) proportion, normalised to Tubulin levels, in *Mfn* dKO stable MEFs cells expressing S-MFN2, L-MFN2 or both isoforms under their endogenous 5’UTR (UTR-MFN2). Quantification was achieved thanks to MFN1 and MFN2 standards produced in E.coli. (n > 4 independent experiments). (B) High-resolution denaturing electrophoresis and western blot of control and *Mfn* dKO stable MEF lines expressing S-MFN2, L-MFN2 or 5’UTR-MFN2. (representative of n > 5 independent experiments). (C) Quantification of mitochondrial morphology in control and *Mfn* dKO stable MEF lines expressing L-MFN2, S-MFN2 or 5’UTR-MFN2. (n ≥ 5 independent experiments, Error bars ± SEM (*p<0.05, **p<0.005, ***p<0.0001)) (D) Representative epifluorescence microscopy of *Mfn* dKO stable MEF lines expressing L-MFN2, S-MFN2 or 5’UTR-MFN2. The impact of the different MFN isoforms expression on mitochondrial morphology (TOM20) and mtDNA (anti-DNA) were analysed using indicated specific antibodies. (scale bar = 10 µm) (E) Quantitative PCR analyses of mtDNA copy number in *Mfn* dKO MEFs, expressing L-MFN2, S-MFN2 or 5’UTR-MFN2. The quantitative PCR was performed using two independent combination of primers targeting nuclear and mitochondrial genomes (left and right bars). (n ≥ 8, Error bars ± SEM ** p<0.01, *** p<0.001, ****p<0.0001) (F) Oxygen consumption rate of *Mfn* dKO MEFs (*Mfn* dKO), expressing L-MFN2, S-MFN2 or 5’UTR-MFN2. The oxygen consumption rate was assessed under endogenous (non-permeabilized), or permeabilized cells in the presence of pyruvate, glutamate malate fuelling the respiratory chain from complex I, under phosphorylating, non-phosphorylating and uncoupled (CCCP titration) conditions. (n ≥ 8, Error bars ± SEM (*p<0.05, **p<0.005, ***p<0.0001)) (G) Cell doubling time of *Mfn* dKO expressing or not the S- or L-MFN2 isoforms in DMEM supplemented with the indicated carbon sources (glucose 25mM; Galactose 10mM in presence of high (10%) or low (3%) fetal bovine serum. (n ≥ 8 independent experiments, Error bars ± SEM (*p<0.05, **p<0.005, ***p<0.0001)).

### S-MFN2 is a key regulator of neutral lipids trafficking and storage

The mitochondrial fusion deficient MEFs (*Mfn1* or *Mfn2* KO MEFs) generated in this study exhibit mitochondrial morphological and dynamic impairments comparable to those observed in *Mfn1* and *Mfn2* KO MEFs previously generated in the Chan Team ^7,42^ (Fig. 3; Supplementary Fig. S5). However, in contrast with these transgenic MEFs models^44,21^, the endoplasmic reticulum (ER) morphology observed in the *Mfn2* KO MEFs we generated was not impaired (Supplementary Fig. S5A; Supplementary Fig. S7A, B). Nevertheless, the mitochondria-ER contacts sites (MERCS) evaluated through classical colocalization^44,21^ or perimeter-based proximity^45^ clearly demonstrated that both the overlapping area and length of the perimeter juxtapositions were modified in response to the mitochondrial morphological aberrations inherent to the loss of MFN2 (Supplementary Fig. S7A-D). Consistent with the findings of Filadi *et al*^45^, our analyses demonstrated that the severity of the MERCS defects is substantially mitigated when the mitochondrial morphological defects of *Mfn2* KO MEFs are rescued by S- or L-MFN2 (Supplementary Fig. S7D). Collectively, our findings demonstrate that both L- and S-MFN2 could restore mitochondrial morphology and, reverse the alterations in ER–mitochondria contact sites.

The increasing number of converging studies supporting MFN2’s role in intracellular lipid trafficking and homeostasis^23,46–48^ prompted us to investigate the mitochondrial lipid profile in newly generated *Mfn2* KO MEFs, as well as the impact of expressing MFN2 mitochondrial isoforms (Fig. 6; Supplementary Fig. S8). First, the analysis of mitochondrial phospholipid composition in *Mfn2* knockout mitochondria remarkably corroborates, the specific signatures of mitochondrial phospholipid deficiencies recently identified in independent fusion-deficient cells (OPA1 or *Mfn1-Mfn2* dKO MEFs)^49^ (Supplementary Fig. S8A, B). Furthermore, we observed that the levels of several neutral lipids detected in the mitochondria fraction were strikingly reshuffled in cells lacking MFN2 (Fig. 6A, B). This accumulation of diacylglycerols, triacylglycerols, acyl-carnitines, and other neutral lipids was efficiently reversed in *Mfn2* KO MEFs stably expressing L-MFN2 or S-MFN2 (Fig. 6A, B). Interestingly, the lipidomic analysis highlighted a specific increase in neutral lipids derived from the mevalonate pathway *i.e.* the esterified cholesterol (cholesteryl esters), and coenzymes Q7 and Q9 only in MEFs expressing S-MFN2 (Fig. 6A, B). This finding was not observed in MEFs expressing L-MFN2, suggesting that the S-MFN2 isoform may have a distinct regulatory role in the trafficking and storage of this neutral lipid subclass. In agreement with a recent study linking MFN2 to elevated cholesteryl ester levels and increased lipid droplet levels^50^, we observed a significant rise in lipid droplet content, quantified by classical Nile Red staining, specifically in cells expressing the murine or human S-MFN2 isoform (Fig. 7; Supplementary Fig. S9). Remarkably, significant changes in cholesterol, coenzyme Q, and neutral lipid storage and trafficking, were specifically linked to the expression of S-MFN2 and did not correlate with the changes measured in mitochondrial morphology or activity (Fig. 4; Fig. 5; Supplementary Fig. S5, Supplementary Fig. S6).

**Figure 6.**
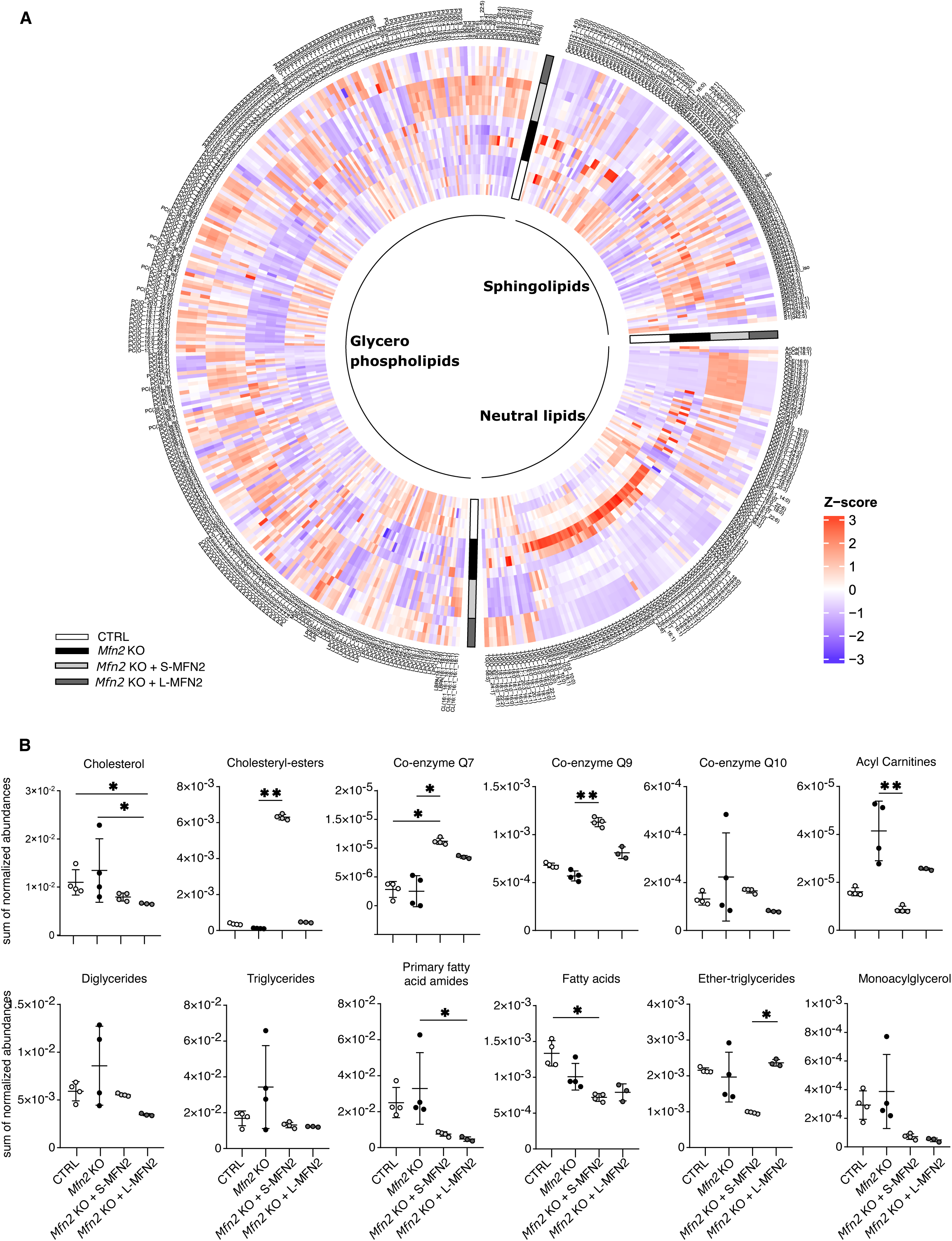
Mitochondrial lipidomic profiling in *Mfn2* KO MEFs expressing S-MFN2 or L-MFN2. (A) Circular heatmap showing the abundances of the identified lipid species across samples. Each concentric circle represents a sample, and color indicates the Z-score of each lipid species across all samples. Red corresponds to a higher abundance relative to the mean abundance of the lipid species across all samples, while blue represents a lower abundance. Lipids are grouped according to their lipid category. (B) Normalized abundance of principal component analysis of neutral lipid profiles obtained by LC-MS (n ≥ 3, mean ± standard deviation). Statistical significance was assessed using the Kruskal-Wallis test (p<0.05) followed by post-hoc Dunn’s test (* p < 0.05, ** p < 0.01).

**Figure 7.**
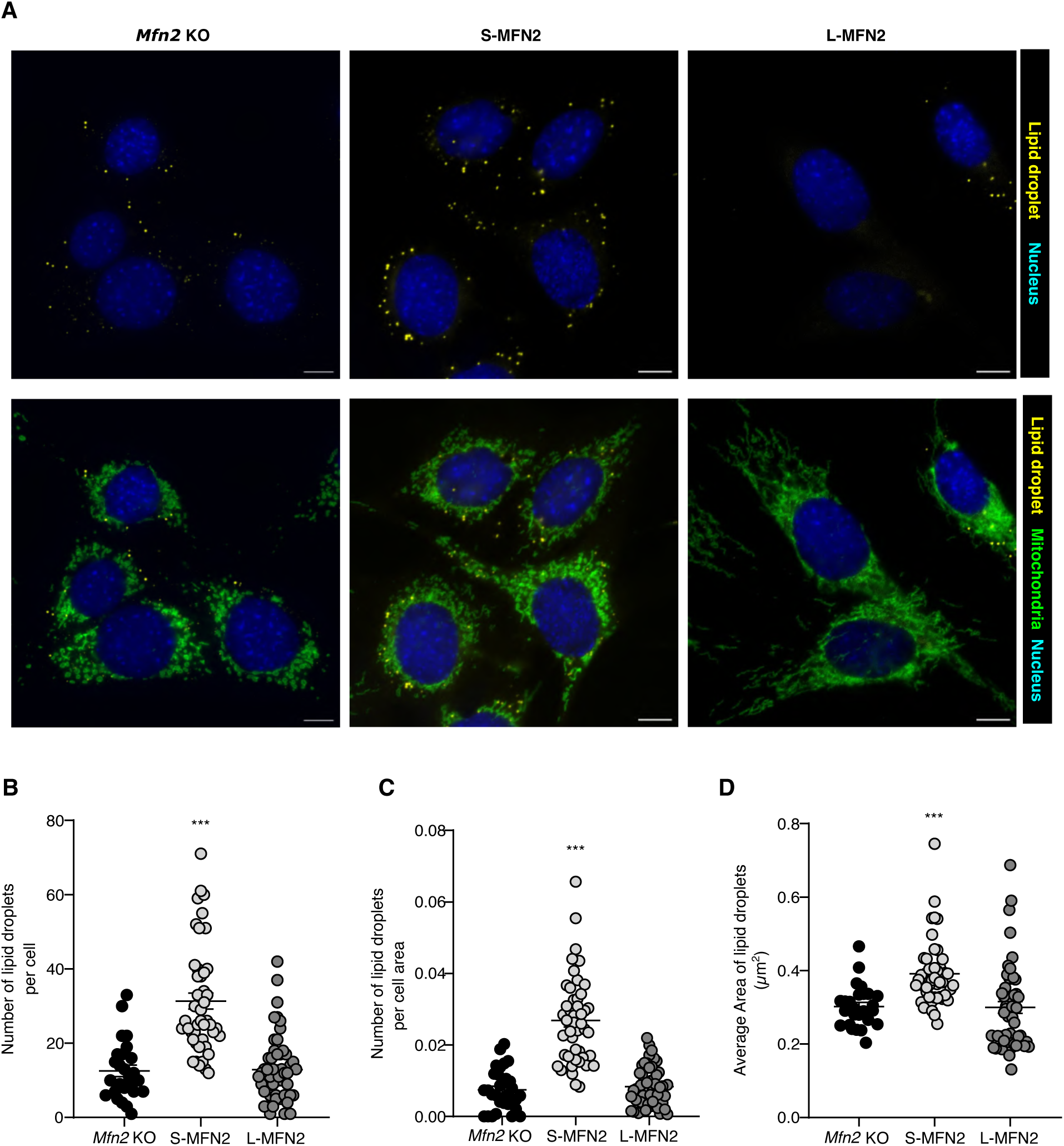
Characterization of lipid droplet levels in *Mfn2* KO MEFs expressing mouse S-MFN2 or L-MFN2. (A) Representative epifluorescence microscopy of *Mfn2* KO stable MEF lines expressing mouse S-MFN2 or L-MFN2, using NileRed as a lipid droplet marker (yellow), DAPI (Blue) and anti-TOM20 as a mitochondrial marker (green) (scale bar = 10 µm). (B) The number of lipid droplets per cell, or cell area (C) and average area (D) were quantified on a single-cell basis in at least 50 cells. The quantification obtained are presented (*n* ≥ 30 cells for *Mfn2* KO, *Mfn2* KO + S-MFN2 and *Mfn2* KO + L-MFN2 MEFs respectively, Error bars indicate ± SEM) and subjected to statistical analysis one-way ANOVA using Dunnett’s multiple comparison test relative to *Mfn2* KO MEFs; *, P < 0.05; **, P < 0.01; ***, P < 0.001).

## DISCUSSION

Mechanisms adjusting the OMM fusion to tissues and cell-types specific needs still remain unknown. Here, we demonstrate that the MFN2 protein doublet resolved by high-resolution denaturing electrophoresis and western blot, are two MFN2 isoforms. As shown in Figure 1A, classical denaturing electrophoresis and western blot analyses does not allow to distinguish the presence of two MFN2 isoforms, which explains why this MFN2 doublet has only been previously noticed a few times, despite the vast bibliography that exists on MFN2 (refs) ^51,52^. Our work unravels that this second MFN2 isoform is generated by an active and highly evolutionary conserved alternative translation ATG site. Moreover, our investigation demonstrates that the tissue-specific expression of the two MFN2 isoforms is orchestrated by splicing of a non-coding exon in the 5’UTR of the gene. Beyond the splicing in 5’UTR, our analysis combining 40S and ribosome foot printing demonstrated that MFN2 5’UTR region encompass several uORFs, using canonical ATG and near cognate CTG, supporting recent findings showing that CTG is frequently used in the context of uORFs^53^. The uORFs are key regulators of gene expression and generally downregulate or suppress translation of the downstream gene^29^. Interestingly, modulating the length of 5’UTR regions through the use of multiple promoters or alternative splicing were previously demonstrated to strongly regulate gene expression ^54^. Two important examples are provided by the breast cancer-associated genes BRCA1 and TGF-ß ^55,56^. Our work demonstrates that splicing of a non-coding 5’exon does not only regulate the expression of the downstream gene but can orchestrate the use of alternative ATGs and regulate the tissue-specific expression of isoforms. We believe that this new genetic regulation will be generalized to other genes presenting aTIS (alternative Translation Initiation Site), as recent genomics and proteomics analyses support that up to 20% of mammalian genes and proteins could be subjected to alternative translation initiation mechanisms ^35,57,58^ and that about 10 to 30% of transcripts exhibit differences in their 5’UTR region ^59^.

The IMM fusion is mainly controlled by the expression of OPA1, which encodes eight isoforms produced by alternative splicing^13,20^. In contrast, neither MFN1 nor MFN2 genes were until recently demonstrated to encode different isoforms. A very recent work has reported the existence of long-range spliced *MFN2* variants^21^. These very short MFN2 variants are deprived of the GTPase domain are therefore not implicated in OMM fusion. Instead, these very low abundant MFN2 isoforms are described to be strictly localized to the endoplasmic reticulum membrane and not involved in mitochondrial fusion. Our work demonstrates that mouse and human MFN2 encode two mitochondrial MFN2 isoforms presenting a 20AA difference in their N-terminus. This 20AA extension differentiating the N-termini of L-MFN2 and S-MFN2, was previously defined as a key sequence differentiating MFN1 from MFN2. In 2019, the publication of Prof. Zorzano’s team devoted great scientific interest in the 20 AA N-terminal extension specifically present in MFN2, showing that this sequence could be specifically involved in phosphatidyl serine (PS) transfer to mitochondria differentiating MFN2 activity from MFN1^47^. Our discovery sheds important new light on this finding, showing that this N-terminal extension is also absent in the most abundant MFN2 isoform found in mouse tissues. Moreover, the existence of a N-terminus shortened isoform (S-MFN2) prompts us to reconsider previous works which have generated and characterized tagged MFN2 proteins. In contrast to the C terminal tag, we now know that the presence of N terminal tag will de facto abrogate the expression of the S-MFN2 isoform.

In line with previous work^38,60,61^, our combined functional analyses indicate that L-MFN2 fusion activity is higher than S-MFN2, but also that the most abundant MFN isoform in brain *i.e.* S-MFN2 has by far the lowest efficiency of fusion. The most complete MFN2 structure was recently resolved^61^ using a N21 truncated version of MFN2, therefore, corresponding to S-MFN2. In line with our observations, functional analyses performed in 2019 demonstrated that the GTPase activity of N20 truncated MFN2 was 13 times lower than MFN1, whereas an 8-fold difference was found by Ishihara in 2004, comparing N terminal-flagged MFN1 and MFN2 (L-MFN2)^60,61^. Combined *in vitro* and *in vivo* experiments will be required to further elucidate how the N21 extension modulates MFN2 fusion activity and potentially confers specialized function to L-MFN2. Our data suggest that the S-MFN2 isoform, despite its reduced ability to rescue mitochondrial morphology and related phenotypes such as mitochondrial genome content and oxidative phosphorylation (OXPHOS) activity, nonetheless plays a distinct role in the regulation of neutral lipid transfer and storage derived from the mevalonate pathway. This lipid signature associated with S-MFN2 corroborates the coenzyme Q deficiencies previously reported in *Mfn2* knockout models of both cardiac tissue and MEFs^23,32^, and shed a new light on numerous studies associating the loss of MFN2 with disruptions in cellular lipid trafficking and storage^46–48,62,63^. The significance of these observations is reinforced by the fact that numerous genes implicated in Charcot-Marie-Tooth (CMT) disease encode key proteins involved in lipid trafficking and by the growing recognition of lipid handling as a critical factor in peripheral neuropathies^64–66^. Future research should determine if the difference in fusion activity and in lipid homeostasis we describe here results from a direct effect between the N-terminal extension and the neighbouring GTPase domain, either by regulating GTPase activity or tethering activities or, more indirectly, by modulating other MFN2-associated functions.

Our work provides the first quantitative analyses of MFN1, L-MFN2, and S-MFN2 levels across tissues and demonstrates that S-MFN2 is the most prevalently expressed MFN isoform in mouse tissue representing about 90% of the MFN expressed in the central nervous system (Fig. 3F, Supplementary Fig. S4D, E; Supplementary Fig.S5D). These observations are in line with a wealth of data generated by different groups demonstrating that MFN1 and MFN2, despite being ubiquitously expressed in all mammalian tissues, present important variability in their proteins levels^37,51,67,68^. Intriguingly, the low fusion efficiency of S-MFN2 seems to correlate with its very high expression levels observed when L-MFN2 and MFN1 are low, as in *Mfn* dKO S-MFN2 cells (Fig. 5A, B) and brain mitochondria (Fig. 3F, Supplementary Fig. S5 D). In contrast, the extremely low abundance of MFN1 in brain, corroborating observations in rat tissues^68^, could provide a very simple and elegant mechanism of the tissue specificity of MFN2 driven neuropathy, and of the specific lethality observed in all generated neuronal *Mfn2* KO mouse models^22,43,69,70^. Conversely, the co-expression of MFN1 and MFN2 in other tissues could explain that loss of MFN2 or MFN1 does not lead to lethal phenotypes in various tissue-specific KO^23,47,71,72^. This extremely high MFN2/MFN1 ratio specifically observed in brain could also explain why MFN1 is dispensable in neurons, in contrast to MFN2^22,43^. The possibility that the low expression of MFN1 in brain is key to explaining the tissue- and MFN isoform-specificities of MFN2 driven neuropathies is also in line with recent observations demonstrating that increasing MFN1 levels in neurons could efficiently rescue adverse phenotypes caused by MFN2 pathogenic mutation^73–76^. Beyond shedding new light to understand both the tissue and isoform (MFN1/MFN2) specificities of the MFN2 driven neuropathies, our work suggests that tissue-specific regulation of the L-MFN2 and S-MFN2 ratio may represent a previously unrecognized mechanism influencing mitochondrial fusion activity.

The previous characterisation by Loiseau and co-workers^77^ of CMT2A patients presenting a point mutation at the methionine 21(M21V) removing the second *MFN2* ATG, demonstrate the crucial role played by the S-MFN2 translation initiation site and strongly enlighten the pathological significance of our discovery. This M21V mutation demonstrates that L-MFN2 cannot complement the absence of S-MFN2 and as a consequence, this mutation causes CMT2A disease. The future development of novel transgenic mouse models expressing S-MFN2 or L-MFN2 will help to understand the respective physiological and functional importance of each MFN2 isoform and provide key information to elucidate the pathogenic mechanisms underpinning CMT2A neuropathies and to open new therapeutic avenues.

## METHODS

### Biological materials

All mice used in this study were on an inbred C57Bl/6N background. Mice were maintained on a standard mouse chow diet and sacrificed, between 20 and 30 weeks of age, by cervical dislocation in strict accordance with the recommendations and guidelines of the Federation of the European Laboratory Animal Science Association, and obtain an authorization from the French Ministry of Agriculture (APAFIS#12648-2017112083056692v7). Mouse embryonic fibroblasts were isolated from homozygous *Mfn1*^Lox/Lox^ or *Mfn2^L^*^ox/Lox^ embryos collected at e13.5 and after few passages were immortalized by transduction with retroviral particles encoding the large T antigen of SV40 (pBABE-puro-SV40 LT, addgene 13970) and then half of the immortalized MEFs generated from one embryo were transduced using Adenovirus vector co-expressing CRE with cytosolic GFP used as reporter. GFP positive *Mfn1* KO and *Mfn2* KO MEFs were FACS sorted and the genetic homogeneity of KO MEFs was then validated by combining previously established genotyping and western blot analyses ^22^. MEFs and Hela cells were grown in DMEM high glucose, 10% FBS, 1% penicillin and streptomycin, and 1% L-Glutamine. The presence of mycoplasma was frequently checked by Dapi staining or PCR tests.

### Monitoring cell proliferation and doubling time

Cells were grown in the following conditions: high-glucose (DMEM supplemented with 10% FBS, 1% penicillin and streptomycin, and 1% L-Glutamine); Low serum (high-glucose DMEM with 3% FBS, 1% penicillin and streptomycin, and 1% L-Glutamine); Galactose (glucose-free DMEM supplemented with 10 mM galactose, 10% FBS, 1% penicillin and streptomycin, and 1% L-Glutamine).

The cells were plated in 6-well plates with 3 mL of the corresponding medium and cultivated for 24, 48, 72, and 96 hours. After each time point, cells were trypsinised, collected, and counted in the presence of trypan blue using the Countess 3 Automated Cell Counter (Thermo Fisher). The doubling time was calculated from the exponential growth phase using standard curve equations.

### Transgenic MEFs expressing 5’UTR variants and mutated ATG, and Hela *MFN2* KD

To generate *Mfn2* KO MEFs expressing 5’UTR containing or not exon 3, 5’UTR from mouse heart (exons 1-2-3) or MEFs (exons 1-2) cDNA were amplified by PCR and then cloned in pQCXIB retroviral vector (Addgene #22800) containing *Mfn2* from Mus.musculus between *Age1-Mfe1* restriction sites. To generate *Mfn2* KO MEFs ΔATG1 or ΔATG2 by mutating ATG1 and ATG2 into ATC and then cloned in a pQCXIB retroviral vector (Addgene #22800). Once validated by sequencing, pQCXIB were transfected in PlatE platinium-E Retroviral Packaging Cell Line, Ecotropic (Cell Biolabs,Inc. cat N°RV-101). Then purified retrovirus was used to transduce *Mfn2* KO MEFs in a DOI allowing to reach about 50% mortality when applying the blasticidin selection. Silencing of *MFN2* expression in Hela cells was achieved using small interfering RNAs ON-TARGETplus human *MFN2* (siRNAs)-*MFN2* siRNAs (L-012961-00-0005) and DharmaFECT1 transfection reagent (T-2001-01) purchased from Horizon Discovery. Hela cells grown to 40-50% confluency were transfected once, which was sufficient to drastically reduce MFN2 protein level assessed by high-resolution western blot.

### Tissues extracts and mitochondria isolation

Quickly after the sacrifice tissues were collected, minced and cleaned with ice cold PBS and snap frozen in liquid nitrogen and then grinded using mortar to obtain tissue extracts powder stored at -80C. To isolate mitochondria, collected tissues were minced and cleaned with ice cold PBS, and suspended in isolation buffer (MIB; 310 mM sucrose, 20 mM Tris-Base, 1 mM EGTA, pH 7.2). Isolation of heart and liver mitochondria was performed by differential centrifugation as previously described ^23,78^. Briefly, pieces of tissues were gently homogenized with few strokes of a Potter S homogenizer (Sartorius) using a loose Teflon pestle, in an ice cold MIB^79^. Mitochondria were isolated by differential centrifugation, after low speed (1,000g, 10 min, 4C) in a swing out rotor, supernatant fraction was subjected to a high speed (7,000g, 10 min, 4C) using a fixed angle rotor. The crude mitochondria pellet obtained was suspended in a minimal amount of MIB. Mitochondrial isolation from MEFs was performed using an Isobiotec cell homogenizer. Briefly, cells were collected using trypsin and the wash with PBS supplemented with soybean trypsin inhibitor (1mg SBTI/mg trypsin) by centrifuging cells at 800g for 10 minutes in swing out rotor. After the final wash of trypsin inhibitor with PBS, cell pellet was suspended in 1.5ml of MIB and subjected to 15-20 strokes of an Isobiotec homogenizer loaded with an 8µm clearance bead. Mitochondria were then isolated by differential centrifugation, after low speed (800g, 10 min, 4C) in a swing out rotor, supernatant fraction was subjected to a high speed (7,000g, 10 min, 4C) using a fixed angle rotor. Protein concentration was measured using Bio-Rad DC Protein Assay kit according to provider instructions.

### High resolution western blot

Tissue extracts (50µg) or isolated mitochondria samples (heart 50µg, skeletal muscle 75µg, liver and kidney 100µg, Brain 5µg) were solubilised in RIPA buffer (NaCl 150 mM, tris-base 50 mM, NP40 1% w/v, SDS 0.1% w/v, deoxycholate 0.5% w/v and EDTA 5 mM, pH 8) supplemented with cOmplete^TM^ protease inhibitor cocktail (Roche) and nuclease (Pierce universal nuclease). After heating samples at 70C for 10 minutes, denatured protein samples were loaded on an Invitrogen^TM^ Bolt^TM^ Bis-Tris 4-12% gradient gel and migration was performed at 100V for 30minutes, 130V for 30 minutes and increased to 150V until the 55 kDa marker reach the bottom of the gel using the MOPS NuPAGE^TM^supplemented with Antioxidant NuPAGE^TM^. After the migration, proteins are blotted to an Amersham^TM^ Protran^TM^ Nitrocellulose premium 0.2µm NC using Invitrogen cassette and BoltTM transfer buffer at 20V for 2 hours. After the transfer, ponceau coloration was systematically performed. Immunodetection was performed by fluorescence using an Amersham^TM^ ImageQuant^TM^ 800. To perform immunodetection membranes were blocked in Rockland’s blocking buffer, then primary antibody was added for overnight incubation, and finally incubated for 1 hour with fluorescent secondary antibody diluted in TBS. The FIJI software was used to determine and analyse densitometric profiles.

The following primary and secondary antibodies were used in this study:

#### Primary antibodies

MFN2 mouse monoclonal antibody clone 6A8 (Abnova H00009927-M01); MFN2 rabbit polyclonal antibody Nter (Abcam 50838), MFN1 rabbit polyclonal antibody (Invitrogen PA5-98602); PMP70 rabbit polyclonal antibody (Abcam ab3421); ERp57/p60 rabbit polyclonal antibody (Proteintech 15967-1-AP); TFAM rabbit polyclonal antibody (Abcam ab131607).

#### Secondary antibodies

ECL Plex goat-mouse IgG, Cy5 (GE Healthcare PA45009); ECL Plex goat-rabbit IgG, Cy5 (GE Healthcare PA45011); CyDye 800 goat-anti-mouse (GE Healthcare 29360788); CyDye 800 goat-anti-rabbit (GE Healthcare 29360791).

### Heart subcellular fractionation

Isolation of heart mitochondria was performed by differential centrifugation as previously described. Briefly, after low speed (1,000g, 10 min, 4C) using a swing out rotor, supernatant fraction was subjected to a high speed (10,000g, 10 min, 4C) using a fixed angle rotor (mitoER fraction). Supernatant fraction was then subjected to very high speed (30,000g, 10min,4°C). We obtained the pellet (ER-perox fraction) and the supernatant fraction (LD fraction). The mito fraction was subjected to a percoll^®^ gradient 17.5% and 20% and centrifuge (10.000g 30min 4°C) to obtained the so-called mito-percoll^®^ fraction and ER-percoll^®^ fraction

### Topology determined by protease shaving

MFN2 isoforms topology analyses were performed on mitochondria isolated from MEF cells expressing L-MFN2 or S-MFN2 and on isolated cardiac mitochondria as they co-express both MFN2 isoforms. Freshly isolated mitochondria (100 µg protein) were incubated in MIB supplemented with trypsin (250µg/ml) at 4C or 37C for up to 30 minutes. To access intermembrane proteins (TIM23) and matrix proteins (TFAM), isolated mitochondria were progressively permeabilized using digitonin concentration ranging from 0.05g/g to 3g/g (g digitonin per g protein). The trypsin reaction was finally quenched by adding a mix of cOmplete^TM^ protease inhibitor cocktail (Roche), PMSF (5 mM) and soybean trypsin inhibitor protease (1mg/ml). After protease quenching, mitochondria were solubilized in RIPA and subjected to low resolution denaturing electrophoresis and western blot.

### MFN1 and MFN2 protein standards purified from *E. coli*

To quantify the proportion of MFN1 and MFN2 proteins in mitochondrial fractions we have produced and purified proteins in *E. coli*. The strain for protein expression and purification was *Escherichia coli* (*E. coli*) BL21(DE3), and all constructs were fully sequenced prior to use. The strain BL21 (DE3) harboring expression plasmid of N terminal His6-tagged Mfn1 or Mfn2 was cultured in Luria Bertani (LB) supplemented with 100 μg/mL ampicillin at 37C with shaking. The culture temperature was then lowered to 20C and after 30 minutes IPTG was added to a final concentration of 1 mM. Cultures were grown for a further 4-6h at 20C and then collected at 5,000 g for 15 min at 4°C. The pellet was solubilized in the lysis buffer (50 mM Tris (pH 8), 250 mM NaCl, 0.5 mM phenylmethylsulfonyl fluoride and protease inhibitor, 0.2% tritonX100) by freeze-thawing and sonication. Mfn1 or Mfn2 was then purified on NiNTA resin. A master mix containing comparable levels of purified MFN1 and MFN2 was prepared and the equal levels of MFN1 and MFN2 were validated by Coomassie and silver staining after denaturing electrophoresis. This MFN1=MFN2 master mix was then diluted and loaded on every high-resolution denaturing electrophoreses to quantify MFN2 and MFN1 signal obtained by western blot to determine the respective proportion of MFN1 and MFN2.

### Alternative splicing detection by PCR

Commercial mouse and human cDNA were purchased from Gentaur-Zyagen: Mouse C57 cDNA, Tissues (heart, skeletal muscles, liver, kidney, brain) Gentaur-Zyagen MD-005-C57, and human heart Tissue cDNA HD-801, human liver cDNA HD-314, human skeletal muscles cDNA HD-102, human brain cDNA HD-201 Gentaur-Zyagen. MEFs cDNA was produced according to the revert aid first strand cDNA synthesis kit from Thermofisher.

The presence of spliced RNA (cDNA) variants was determined by designing a combination of 7 forward and 6 reverse primers covering all exons of mouse or human mRNAs. The primers used are as follow:

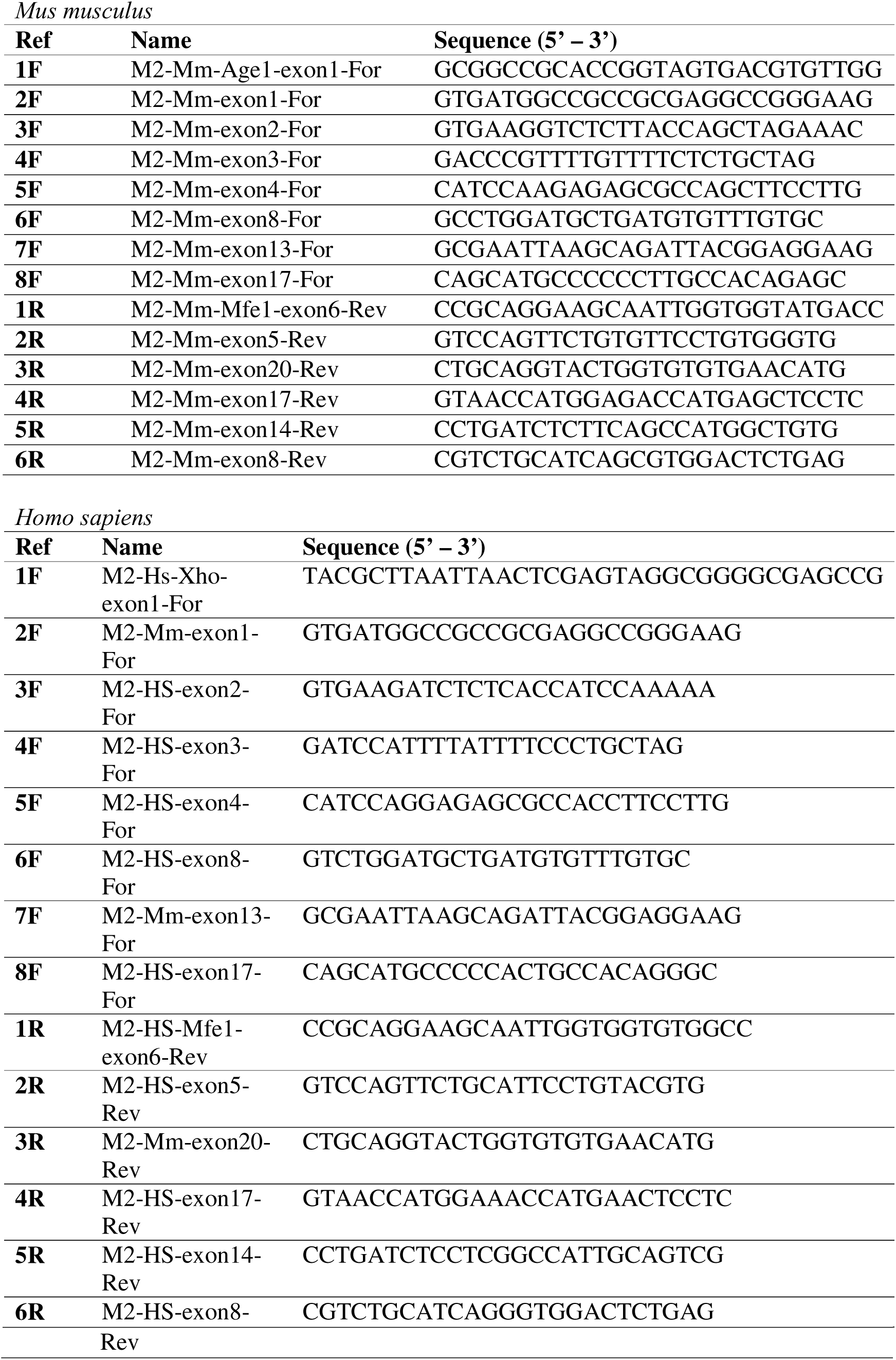

Every truncated version identified in mouse or human as a spliced variant has been validated by sequencing. To this end, PCR products were purified with GeneJet extraction Kit (Thermo scientific K0691) and further amplified using Mix2Seq Kit and service (Eurofins/genomics).

### mtDNA/nDNA ratio determined by quantitative PCR

Total DNA was extracted from the different immortalized MEF cell lines at 70% confluence, using the DNeasy Tissue and Blood Kit by Qiagen (Cat.No. 69504) following the manufacturer’s instructions. Total DNA quantification was achieved using the Helixyte Green^TM^ dsDNA Quantification Kit Green Fluorescence by AAT Bioquest (Cat.No. 17651). Then, 10 ng of total extracted DNA was used to achieve a quantitative real-time PCR using the 2X GoTaq qPCR MasterMix by Promega (Cat. No. A6001) along with 2 μM of the reverse and forward primers amplifying mitochondrial DNA and the reverse and forward primers amplifying nuclear DNA. Cycling conditions were as follows: 95°C for 3 min for one cycle, then a 3-step protocol, 95°C for 30 seconds, 64°C for 30 seconds and 72°C for 1 minute, for 40 cycles. Using a calibration range method for each primer, the abundance of mitochondrial DNA was determined as a ratio generated by normalizing the value obtained from the mitochondrial probe to the value from the nuclear probe which is subsequently followed by a normalization to the control.

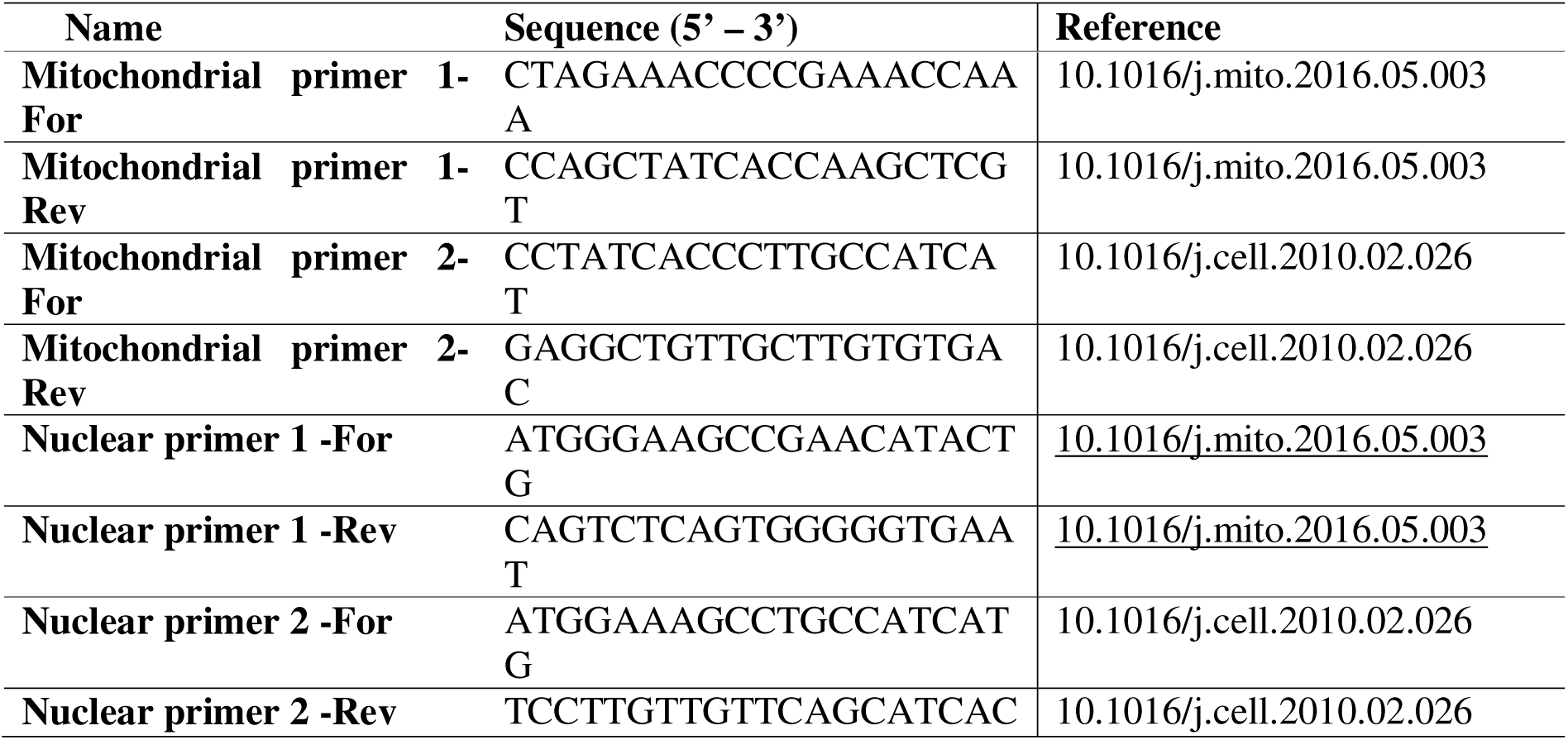

### High-resolution oxygen consumption

High-resolution oxygen consumption rate of digitonin-permeabilized cells was performed as described previously^32^ at 37°C with slight modifications. 50µL of cells (∼ 1 mg total cellular protein) were diluted in 0.5 mL of respiratory buffer (120 mM sucrose, 50 mM KCl, 20 mM Tris-Base, 4 mM KH_2_PO_4_, 2 mM MgCl_2_, 1 mM EGTA, pH 7.2 at RT) within the chamber of an Oxygraph-2K (Oroboros). Following whole cell respiration stabilization, cells were permeabilized by successive addition of digitonin concentrated at 0.2 mg/ml. Optimum cells permeabilization was obtained with a digitonin range from 2.4 to 4.8 µg/ml depending of the cell lines studied. The oxygen consumption rate under phosphorylating condition was assessed using either NADH-linked substrates (10 mM glutamate, 10 mM pyruvate and 5 mM malate), or FADH-linked substrates (10 mM succinate plus 5 mM glycerol-3-phosphate in presence of 8 nmol/L of rotenone) in the presence of 2 mM ADP. The non-phosphorylating state was obtained after ATP synthesis inhibition using 30 ng/ml oligomycin. Maximal mitochondrial respiration was obtained following successive addition of CCCP, steps of 50 nmol/L. Oxygen consumption was expressed in nat O/min/mg proteins.

### Bioinformatic analyses of mouse tissue-specific 5’UTR splicing

Raw sequencing data from the *Tabula muris* bulk sequencing project^24^ were downloaded from the GEO website (accession number GSE132040). For each analyzed tissue, we used transcriptome data from 2 males and 2 females of 9 months. For differential splicing analysis, we used Spladder^80^, comparing each tissue against muscle.

### Fluorescent microscopy – image aquisition

Immunofluorescence microscopy was essentially performed as described previously^81^. Cells plated onto glass coverslips were fixed in 3.2% paraformaldehyde for 20 min at room temperature, then washed once with PBS before being permeabilized using PBS-Triton X 0.1% (PBST) solution for 5 min. For both endoplasmic reticulum and peroxisome staining’s, the coverslips were incubated for 20 min in an 8M urea solution, then washed with PBST once before incubation for 1h30 min at room temperature with the following primary antibodies prepared in PBS: rabbit polyclonal IgG anti-Calnexin (abcam, ab22595, dilution of 1/200) as an endoplasmic reticulum marker and rabbit polyclonal IgG anti-PMP70 (abcam, ab3421, dilution of 1/400) as a peroxisome marker. Incubation with the following secondary antibody goat anti-rabbit Alexa 555 (dilution of 1/500) was then achieved for 45 min at room temperature. Finally, the coverslips were washed with PBST, then PBS, and distilled water before being mounting with DAPI-containing MOWIOL. For DNA and mitochondrial staining, coverslips were incubated for 30 min in a 10% BSA solution, then washed with PBST once before incubation for 1h30 min at room temperature with the following primary antibodies prepared in 3% BSA solution: mouse IgM anti-DNA (Progen, 61014, dilution of 1/250) and rabbit anti-TOM20 (Santa Cruz, sc-11415, dilution of 1/400). Incubation with the following secondary antibodies goat anti-mouse IgG(H+L) Alexa Fluor^TM^ Plus 488 (Invitrogen, A32723 dilution of 1/500) and goat anti-rabbit IgG (H+L) Alexa Fluor^TM^ Plus 555 (Invitrogen, A32732, dilution of 1/500) was then achieved for 45 min at room temperature. Finally, the coverslips were washed with PBST, then PBS, and distilled water before being mounting with DAPI-containing MOWIOL. Fluorescence images were acquired using an inverted Microscope Olympus (Olympus IX81).

To estimate mitochondrial-endoplasmic reticulum contacts, coverslips were incubated for 30 min in a 10% BSA solution, then washed with PBST once before incubation for 1h30 min at room temperature with the following primary antibodies prepared in 3% BSA solution: mouse anti-ATPase IF1 (abcam, ab110277, dilution of 1/100) and rabbit anti-calnexin (Santa Cruz, abcam, ab22595, dilution of 1/200). Incubation with the following secondary antibodies goat anti-rabbit IgG(H+L) Alexa Fluor^TM^ Plus 488 (Invitrogen, A32723 dilution of 1/500) and goat anti-mouse IgG (H+L) Alexa Fluor^TM^ Plus 555 (Invitrogen, A32732, dilution of 1/500) was then completed for 40min at room temperature. Finally, the coverslips were washed with PBST, then PBS, and distilled water before being mounted with DAPI-containing MOWIOL. Fluorescence images were acquired using an inverted ECLIPSE Ti2 microscope.

To visualize and quantify lipid droplets, coverslips were incubated for 30 min in a 10% BSA solution, then washed with PBST once before incubation for 1h30 min at room temperature with rabbit anti-TOM20 (Santa Cruz, sc-11415, dilution of 1/400) prepared in 3% BSA solution which was followed by a 40min incubation at room temperature with the secondary antibody goat anti-rabbit IgG (H+L) Alexa FluorTM Plus 488 (Invitrogen, A32732, dilution of 1/500). The coverslips were then incubated with Nile Red (Invitrogen, N1142, 100ug/mL) for 30min at room temperature and were washed with PBST, then PBS, and distilled water before being mounting with DAPI-containing MOWIOL. Fluorescence images were acquired using an inverted ECLIPSE Ti2 microscope.

To correlate mitochondrial morphology and mitochondrial DNA to the level of MFN2 protein, coverslips were incubated for 30 min in a 10% BSA solution, then washed with PBST once before incubation for 1h30 min at room temperature with the following primary antibodies prepared in 3% BSA solution: rabbit anti-MFN2 D1E9 (Cell Signaling Technology, 11925, dilution 1/100), rabbit anti-MFN2 D2D10 (Cell Signaling Technology, 9482, dilution 1/100), mouse ATPase IF1 (abcam, ab110277, dilution of 1/100) and mouse anti-DNA (Progen, 61014, dilution of 1/200). Incubation with the following secondary antibodies goat anti-rabbit IgG(H+L) Alexa Fluor^TM^ Plus 488 (Invitrogen, A32723 dilution of 1/500) and goat anti-mouse IgG (H+L) Alexa Fluor^TM^ Plus 555 (Invitrogen, A32732, dilution of 1/500) was then completed for 40min at room temperature followed by a 30min incubation with Alexa Fluor™ 647 phalloidin (Invitrogen, A22287, 1/1000). The coverslips were washed with PBST, then PBS, and distilled water before being mounting with DAPI-containing MOWIOL. Fluorescence images were acquired using an inverted ECLIPSE Ti2 microscope.

### Fluorescent microscopy – image analysis

For the single cell analysis of mitochondrial network, images were pre-processed and segmented, before proceeding to the 2D analysis using Mitochondria Analyzer, which provides mitochondrial count, area, perimeter, branch number and length, and junctions number, among other available parameters. For the mitochondrial DNA assessment, the obtained images were thresholded on a single cell basis. Following segmentation and binarization, the nuclear area was manually removed/cropped from the image before analysis using the particle analyzer command, which provides particle counts, area, and size among other parameters. To obtain the MFN2 expression level for each cell, all images were acquired using the same exposure time. In order to focus on the signal intensity of MFN2 solely, thresholding of the images was completed before measuring signal intensity limited to the thresholded area.

For lipid droplet analysis, object detection was achieved using the StarDist 2D plugin, the versatile model (fluorescent nuclei) being chosen as neural network prediction. The obtained identified ROI were then restored onto the original image to analyze the total count and size of each droplet.

For the mitochondria-endoplasmic reticulum contact analysis, mitochondrial area and endoplasmic reticulum area were calculated after auto-local threshold. Pearson’s and Mander’s colocalization indexes were obtained using the BIOP JACoP. To trace the profile of the mitochondrial and the endoplasmic reticulum, the images were transformed into binary images which were then run through the IsoPhotContour2 plugin by ImageJ. The images of the mitochondrial profile were then merged with the ER-mitochondria colocalization pixels before extracting colocalization parameters such as number and perimeter of the identified contacts.

### Expansion microscopy

Ultrastructure expansion microscopy (U-ExM) was performed essentially as described ^39^. M*fn2* KO MEFs expressing MFN2 isoforms carrying fluorescent protein tags (MFN2S-mCherry and MFN2L-EGFP) were fixed with PBS containing 3% paraformaldehyde and 0.1% glutaraldehyde (20 minutes, room temperature). Fixed cells were incubated for 3h at 37°C in PBS containing 0.7% formaldehyde and 1% acrylamide, washed and subjected to gelation. The acrylamide gel contained 10% acrylamide, 0.025% bis-acrylamide and 19% sodium acrylate; lowering the bis-acrylamide concentration of the original U-ExM recipe (0.1%) increased the expansion factor from roughly 4X to roughly 6X. Gel hydration, denaturation and expansion were performed as described. For labelling with antibodies against EGFP, mCherry and/or VDAC, gels were transferred to PBS and incubated over night at 37°C with primary antibodies. Gels were washed three times for 10 min and incubated for 2 hours at 37°C with secondary antibodies. Gels were washed extensively with PBS and fully expanded in H_2_O. The size of the expanded gel (∼68-72 mm) allowed to determine the overall expansion factor (5,7-6X). Antibodies were diluted in the washing solution (PBS+0.1% Tween-20). Primary antibodies: anti-EGFP serum generated in rabbits immunized with purified EGFP (agro-bio.stago, France), goat anti-mCherry (biorbyt orb11618) and mouse anti-VDAC (abcam, ab14734). Secondary antibodies: donkey anti rabbit IgG-AlexaFluor 488+ (Thermofisher Scientific # A32790), donkey anti goat IgG-CF555, (biotium 20039) and donkey-anti-mouse CF640R (biotium 20177). Gels were immobilized on glass bottom dishes (ibidi, 81158) treated with 0.1% poly-lysine and visualized in an inverted Olympus IX81 microscope. Images were treated with ImageJ.

### Antibodies list

| *Primary antibodies* |  |  |  |
| --- | --- | --- | --- |
| Antibody designation | Supplier | Catalog number | Source |
| Mfn2cTer clone6A8 | Abnova | H00009927-M01 | Mouse |
| Mfn2 nTer | Abcam | Ab50838 | Rabbit |
| Mfn1 | Invitrogen | PA5-98602 | Rabbit |
| PMP70 | Abcam | Ab3421 | Rabbit |
| ERp57/ERp60 | Proteintech | 15967-1-AP | Rabbit |
| TFAM | Abcam | Ab131607 | Rabbit |
| Tom70 | Invitrogen | PA1-46188 | Rabbit |
| Tom20(FL-145) | SantaCruz | sc-11415 | Rabbit |
| Tom22 | Abcam | ab57523 | Mouse |
| RRBP1 | Proteintech | 22015-1-AP | Rabbit |
| SERCA2ATPase | Invitrogen | MA3-919 | Mouse |
| Tim23 | BD Biosciences | 611222 | Mouse |
| Calnexin | Abcam | Ab22595 | Rabbit |
| VDAC | Abcam | Ab14734 | Mouse |
| Tubulin(DM1A) | Sigma | T9026 | Mouse |
| DNA,AC-30-10 | Progen | 690014 | Mouse |
| EGFP | agro-bio.stago | Custom made | Rabbit |
| mCherry | biorbyt | orb11618 | Goat |

| *Secondary antibodies* |  |  |
| --- | --- | --- |
| **Antibody designation** | **Supplier** | **Catalog number** |
| ECL Plex goat-mouse IgG, Cy5 | GE Healthcare | PA45009 |
| ECL Plex goat-rabbit IgG, Cy5 | GE Healthcare | PA45011 |
| CyDye 800 goat-anti-mouse | GE Healthcare | 29360788 |
| CyDye 800 goat-anti-rabbit | GE Healthcare | 29360791 |
| Peroxidase-AffiniPure Goat Anti-Mouse | Jackson ImmunoResearch | 115-035-062 |
| Peroxidase AffiniPure Goat Anti-Rabbit | Jackson ImmunoResearch | 111-035-045 |
| Donkey anti goat IgG-CF555 | biotium | 20039 |
| Donkey-anti-mouse CF640R | biotium | 20177 |
| Alexa fluor plus 488 goat-anti-mouse | Invitrogen | A32723 |
| Alexa fluor plus 555 goat-anti-rabbit | Invitrogen | A32732 |

### Ribosome profiling data analysis

40S-seq and GTI-seq data from HEK293T cells were downloaded from GEO (accession number GSE174329)^26^. For 40S-seq data analysis, SRR14510016, SRR14510017, SRR14510018, and SRR14510019 data sets were combined. For GTI-seq data analysis, SRR14510026 and SRR14510027 data sets were combined. First, read lengths were filtered (20-60 nucleotides) using Cutadapt^82^. Next, reads were aligned to human MFN2 transcript variant X3 (NCBI Reference Sequence: XM_005263543.4), variant 1 (NCBI Reference Sequence: NM_014874.4), and variant 2 (NCBI Reference Sequence: NM_001127660.2) or to human MFN1 transcript (NCBI Reference Sequence: NM_033540.3) using Bowtie2 end-to-end alignment ^83^. Aggregated read counts were obtained via Plastid applying centre-weighted read mapping (DOI: 10.1186/s12864-016-3278-x).

Ribo-seq data from mouse heart were downloaded from European Nucleotide Archive (ENA) (accession number PRJEB29208; run accession: ERR3367791)^84^. First, adaptor sequences (AGATCGGAAGAGCACACGTCT) were trimmed and read lengths were filtered (20-36 nucleotides) using Cutadapt^82^. Next, reads were aligned to mouse MFN2 transcript variant 1 (NCBI Reference Sequence: NM_001285920.1), and variant 2 (NCBI Reference Sequence: NM_133201.3), or to mouse MFN1 transcript (NCBI Reference Sequence: NM_024200.5) using Bowtie2 end-to-end alignment^83^Aggregated read counts were obtained via Plastid applying centre-weighted read mapping^84^.

### Lipidomics analysis and data analysis

Mitochondrial isolation from MEFs (Control, *Mfn2* KO, S-MFN2 and L-MFN2) was performed using an isobiotec cell homogenizer as previously described. Mitochondrial lipid extraction was performed as follows: 300 µL of isopropanol were added to a dry pellet of 100 µg of isolated mitochondria, vortexed thoroughly and sonicated for 10 minutes. The mix was then centrifugated (15,000 g, 10 min, 4°C) and 250 µL of the supernatant was recovered, put in a glass vial for solvent evaporation. The resulting residue was reconstituted with 100 µL of acetonitrile/isopropanol/water (6:3:1).

LC-HRMS analysis was performed using a UHPLC Ultimate 3000 system (Dionex) coupled to a Q-Exactive mass spectrometer (Thermo Fisher Scientific) operated in positive and negative electrospray ionization modes. Chromatography was carried out with a ACQUITY UPLC BEH C18 column (100 mm × 2.10 mm x 1.7 µm, 130Å) (Waters) heated at 55°C. The solvent system comprised mobile phase A [0.1% formic acid in water], and mobile phase B [0.1% formic acid in acetonitrile]. Chromatographic separation was performed at a flow rate of 0.4 mL/min as follows: 0 to 2.7 min, 90 %A ; from 2.7 to 2.8 min, 55 %A ; from 2.8 to 8 min, 47 %A ; from 8 to 8.1 min, 40% A ; from 8.1 to 11.5 min, 20 %A ; from 12 to 13.2 min, 0 %A; from 13.2 to 15 min, 90 %A.

The autosampler (Ultimate WPS-3000 UHPLC system, Dionex) was set at 4°C and the injection volume at 5 μL for each sample. The heated ESI source parameters were a spray voltage of 3.5 kV, capillary temperature of 350°C, heater temperature of 250°C, sheath gas flow of 35 arbitrary units (AU), auxiliary gas flow of 10 AU, spare gas flow of 1 AU, and tube lens voltage of 60 V for C18. During the full-scan acquisition, which ranged from 250 to 1600 m/z, the instrument operated at 70,000 resolution, with an automatic gain control target of 1 × 106 charges and a maximum injection time of 250 ms.

The instrumental stability was evaluated by multiple injections (n = 5) of a quality control (QC) sample obtained from a pool of 10 μL of all samples analysed. This QC sample was injected at the beginning of the analysis, in between the sample injections, and at the end of the run.

Following analysis, data were processed using workflow4metabolomics (23.0 release)^85^. The level of each feature was expressed as a normalized abundance, calculated by dividing its intensity by the sum of all feature intensities in that sample for each modality. Lipid annotation was performed using LipidSearch 5.1 (ThermoScientific) on MS2 data acquired from the QC sample in both positive and negative ionization modes, following current lipid nomenclature guidelines^86,87^. Annotations were manually reviewed but remain putative. Features with a coefficient of variation greater than 30% in the QC samples were excluded from further analysis. Positive and negative processed datasets were finally combined and identified lipids were manually sorted to avoid redundancy.

### Statistics

For the circular heatmap representation, Z-score normalization was applied to the normalized abundance of each lipid across all samples. The Z-score for each lipid in a given sample was calculated as: Z = (normalized abundance of the lipid in the sample - mean normalized abundance of that lipid across all samples) / standard deviation of that lipid’s normalized abundance across all samples. For lipid class analysis, the normalized abundances of all lipids within a given class were summed per sample. For multiple group comparisons, Kruskal-Wallis test followed by Dunn’s post-hoc test was used. Data are presented as mean ± standard deviation. Principal component analysis was performed on the normalized abundances of the identified lipid species.

## DATA AVAILABILITY

The authors declare that the main data supporting the findings of this study are available within the article and its Supplementary Information files. Extra data are available from the corresponding author upon request.

## ACKNOWLEDGEMENTS

We would like to thanks the Animalerie A2, led by Julien Izotte and Benoit Rousseau for housing the different *Mfn* mouse colonies (the service commun des animaleries, Animalerie A2 – University of Bordeaux). We would also like to thanks the Plateforme de Vectorologie of Bordeaux University, led by Véronique Guyonnet Dupérat and Violaine Moreau for providing Adenovirus CRE-GFP to generate MEF lines. This work was supported by the “Plateforme de Métabolomique et d’Analyses Chimiques, US61 ASB, Université de Tours, CHRU Tours, Inserm, Tours, France” and by the MetaboHUB infrastructure funded by the Agence Nationale de la Recherche under the France 2030 program (MetaboHUB ANR-11-INBS-0010 ; MetEx+ ANR-21-ESRE-0035; MetaboHUB (JVCE) ANR-24-INBS-0012). We thank Thomas Roberts for pBABE-puro SV40 LT (addgene #13970) and Eric Campeau for pQCXIB (addgene #22800).

## FUNDINGS

This work was supported by the ANR (ANR-16-CE14-0013 and ANR-22-CE14-0040), Bordeaux University (GPR Light) and region Nouvelle Aquitaine (MetabOptic CRNA ESR2023). The funders had no role in study design, data collection and analysis, decision to publish, or preparation of the manuscript.

## AUTHORS CONTRIBUTIONS

A.M. conceptualized research goals and experiments supervise and together with C.D contributed to develop most of the model and experimental procedure use in this work; C.D planned, performed and analyzed results from the majority of experiments, also developed most of the MEFs models and greatly contributed to the conceptual discovery; C.B and C.E performed mtDNA quantification and C.B performed microscopy analyses using conventional epifluorescent microscope and developed the single cell analyses; C.E performed several western blot and together with M.P performed the lipidomic analyses; P.P and G.D. performed bioenergetic analyses ; M.P. performed lipid extractions, lipid identification, analyzed and interpreted the lipidomic data, prepared the lipidomic illustrations. A.L. performed the LC-MS/MS experiments; M.R and J.D performed expansion microscopy analyses and supervised the microscopy analyses ; C.D, R.B., E.C and T.M, contributed to generate MEFs models and developed high resolution electrophoreses. A.K and J.R. performed bioinformatic analyses of ribosome footprinting dataset; B.H. performed bioinformatic analyses to identify, validate and correlate 5’UTR splicing across mouse tissues. AM wrote the manuscript and generated figures with input of all authors.

## COMPETING INTERESTS

The authors declare no conflict of interest.

---

**Figure S1.**
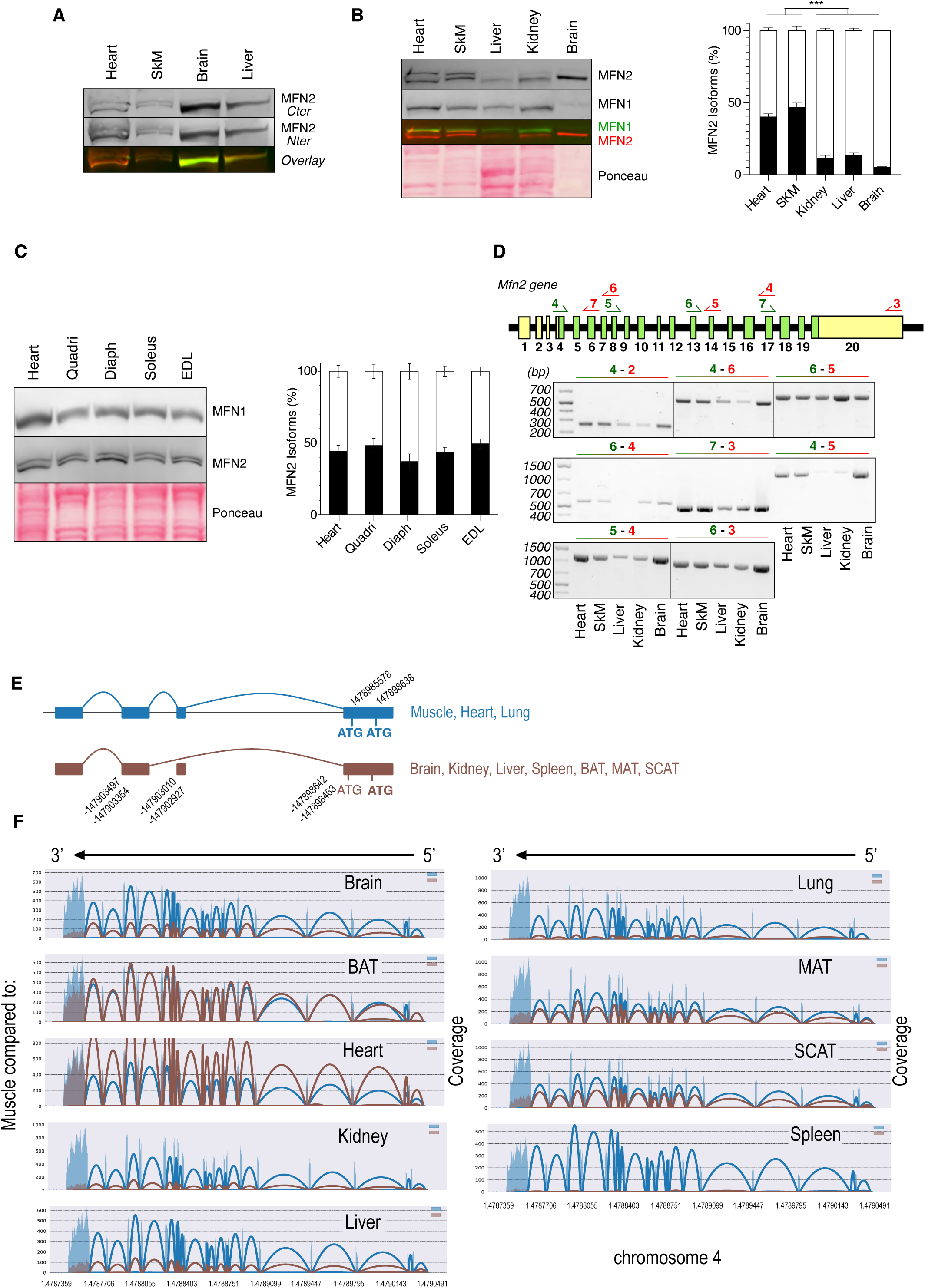
High resolution denaturing electrophoresis and western blot unravels MFN2 protein doublet. (A) High-resolution denaturing electrophoresis and western blot, performed on total protein extracts from indicated tissues, using two different MFN2 antibodies targeting Cter or Nter epitopes. (representative of n ≥ 4 independent experiments). (B) High-resolution denaturing electrophoresis and western blot, performed on mitochondrial protein extracts from indicated tissues, using two different MFN1 and MFN2 antibodies (n=6 mitochondria isolations). On the right, Proportion of upper (black bars) and lower (white bars) proteins present in the MFN2 doublet, determined by densitometric analyses of high-resolution western blot analyses performed on purified mitochondrial protein extracts (n ≥ 6 independent experiments). Error bars ± SEM (***p<0.0001). (C) High-resolution denaturing electrophoresis and western blot, performed on mitochondrial protein extracts from indicated muscle tissues, using two different MFN1 and MFN2 antibodies (n=6 mitochondria isolations). Proportion of upper (black bars) and lower (white bars) proteins present in the MFN2 doublet, determined by densitometric analyses of high-resolution western blot analyses performed on purified mitochondrial protein extracts (n ≥ 6 independent experiments). Error bars ± SEM (***p<0.0001). (D) PCR detection of *Mfn2* splice variants performed within the coding sequence of cDNA of the indicated mouse tissues (under each PCR products). (top) Forward primers (green half arrows and numbers) and reverse primers (red half arrows and numbers) are positioned on *Mfn2* ORF. Non coding and coding regions respectively appear in yellow and green. The couple of primers used for the reaction is indicated on top of each reaction products. (representative of n ≥ 3 independent experiments) (E) Schematic view summarizing bioinformatic analysis of Tabula muris sequencing data, identity of tissues presenting or not 5’UTR exon splicing are listed as well as the genomic positions of the different exons and ATG. (F) Representation of the *Mfn2* transcriptomic profile determined in the indicated mouse tissues and present in the Tabula muris sequencing project and analysed with Spladder bioinformatic tools.

**Figure S2.**
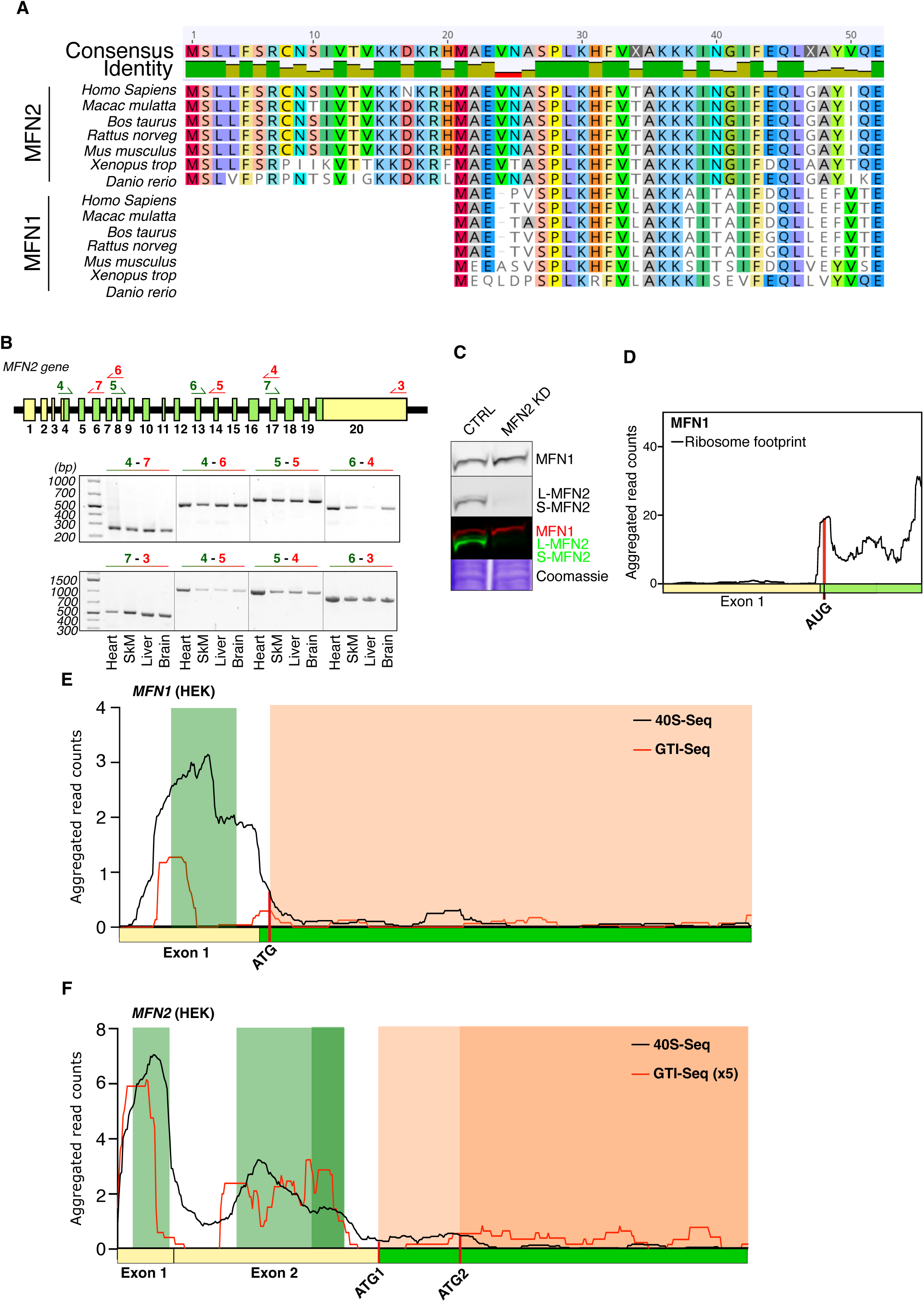
Alternative ATG is evolutionary conserved. (A) Protein pairwise sequence alignment of MFN1 and MFN2, across model organisms performed with Geneious software. (B) PCR detection of *MFN2* splice variants performed within the coding sequence of cDNA purified from the indicated human tissues (under each PCR products). (top) Forward primers (green numbers and half arrow) and reverse primers (red numbers and half arrow) are positioned on *MFN2* ORF. Non coding and coding regions respectively appear in yellow and green. The couple of primer used for the reaction is indicated on top of each PCR reaction products. (representative of n ≥ 3 independent experiments) (C) High-resolution denaturing electrophoresis and western blot, performed on total extract from Hela control and *MFN2* KD cells (representative of n=3 independent experiments). (D) Alignment of existing ribo-seq data from mouse heart tissue to MFN1 transcript^88^. Region comprising 1-389 nucleotides is shown. Coding and non-coding region are coloured in green and yellow respectively. Location of translation initiation codon ATG is presented with a red line. (E and F) Alignment of existing TISCA data from HEK293T cells to *MFN1* and *MFN2* transcripts^26^. Regions comprising 1-459 nucleotides are shown for both transcripts. Start sites are indicated by a combined presence of a GTI-seq peak (approximately +12 nucleotides from 5’-end of GTI-seq reads) and a decrease of 40S-seq reads at approximately 24 nucleotides upstream of the GTI-seq peak. The presence of sequence-compatible upstream ORF (uORF) is symbolized with a green background and coding ORF by an orange background. The bottom line in the x-axis is the sequence composed by non-coding (yellow) and coding (green) regions. Translation initiation codons are highlighted by red lines.

**Figure S3.**
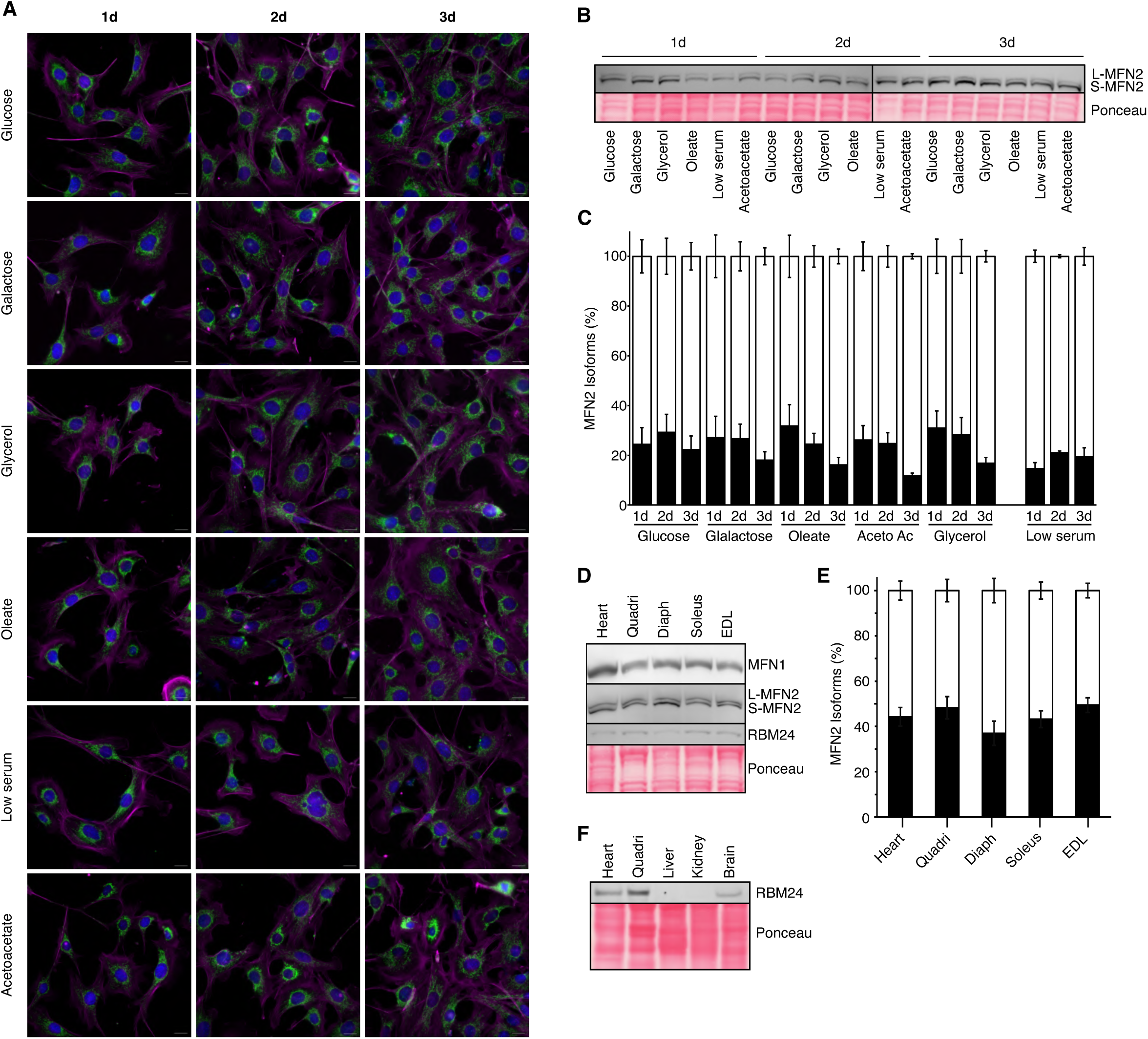
The MFN2 isoforms expression remain unchanged under various metabolic stress conditions. (A) Representative epifluorescence microscopy of control MEFs cultivated during 1, 2 or 3 days in DMEM medium supplemented with glucose, galactose (10mM), glycerol (10mM), Oleate (500µM complexed with BSA according to a 3:1 ratio), acetoacetate (10mM). The mitochondrial morphology was followed using anti-TOM20 as a mitochondrial marker (green), staining against phalloidin as a cytoskeleton marker (magenta) and DAPI (scale bar = 10 µm). (representative n = 3 independent experiments) (B) High-resolution denaturing electrophoresis and western blot of MFN2 performed with total protein extracts from control MEFs cultivated during 1, 2 or 3 days in DMEM medium supplemented with indicated metabolic conditions (representative n = 4 independent experiments). (C) Proportion of S-MFN2 (white bars) and L-MFN2 (black bars) present in the MFN2 doublet, determined by densitometric analyses post high-resolution denaturing electrophoresis and western blot analyses performed on total protein extracts from harvested cells (n = 4 independent experiments, Error bars ± SEM). (D) High-resolution denaturing electrophoresis and western blot of MFN1, MFN2 and RBM24 performed with total protein extracts from indicated mouse muscles tissues presenting divergent energy metabolisms (representative n = 4 independent experiments). (E) Proportion of S-MFN2 (white bars) and L-MFN2 (black bars) present in the MFN2 doublet, determined by densitometric analyses of western blot analyses performed on total protein extracts from the indicated mouse muscles tissues (n = 4 independent experiments; Error bars ± SEM). (F) RBM24 proteins levels detected by denaturing electrophoresis and western blot performed with total protein extracts from the indicated mouse tissues (n = 4 per tissue).

**Figure S4.**
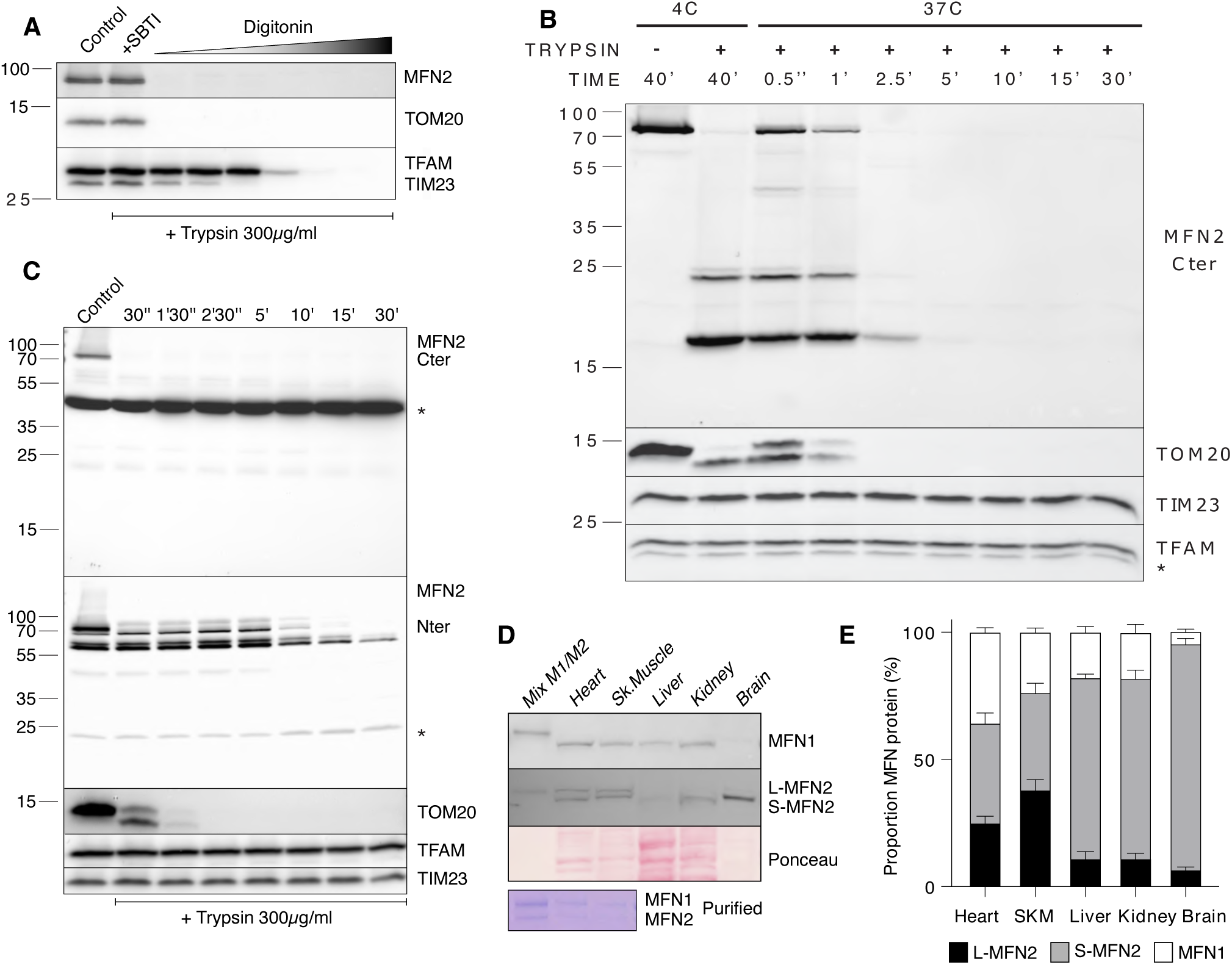
S-MFN2 exhibit canonical MFN2 topology and is the most abundant MFN2 isoform in mouse tissues. (A) Trypsin protease shaving analyses performed on isolated heart mitochondria showing that MFN2 isoforms behave as TOM20. The range of digitonin to protein ratio used allows for efficient discrimination of extramitochondrial, intermembrane and intramitochondrial compartments finely controlling the OMM and IMM permeabilization and protease accessibility. The second lane is a control showing that the trypsin inhibitor used can efficiently prevent the action of trypsin. (representative of n = 4 independent experiments). (B) Trypsin protease shaving analyses performed on isolated mitochondria from control MEFs. The Trypsin protease shaving was performed on non-permeabilized mitochondria for 40 minutes at 4C or followed over 30 minutes kinetic at 37C. (representative of n = 3 experiments) (C) Trypsin protease shaving analyses performed on isolated heart mitochondria from control mice. The Trypsin protease shaving was performed on non-permeabilized mitochondria and followed over 30 minutes kinetic at 37C. (representative of n = 3 experiments, * indicates aspecificity) (D) High-resolution MFN2 denaturing electrophoresis and western blot performed on isolated mitochondria from indicated mouse tissues. The lower panel represents Coomassie staining performed after denaturing electrophoresis performed with MFN1 and MFN2 produced in E.coli and used in high-resolution western blot for quantification (representative of n = 4 experiments). (E) Quantification of MFN proteins using quantitative western blot analysis. The proportion of the between MFN1, L-MFN2 and S-MFN2, across isolated mitochondria from indicated mouse tissues, was quantified using MFN1 and MFN2 standards produced in E.Coli. (n = 8 mitochondria samples).

**Figure S5.**
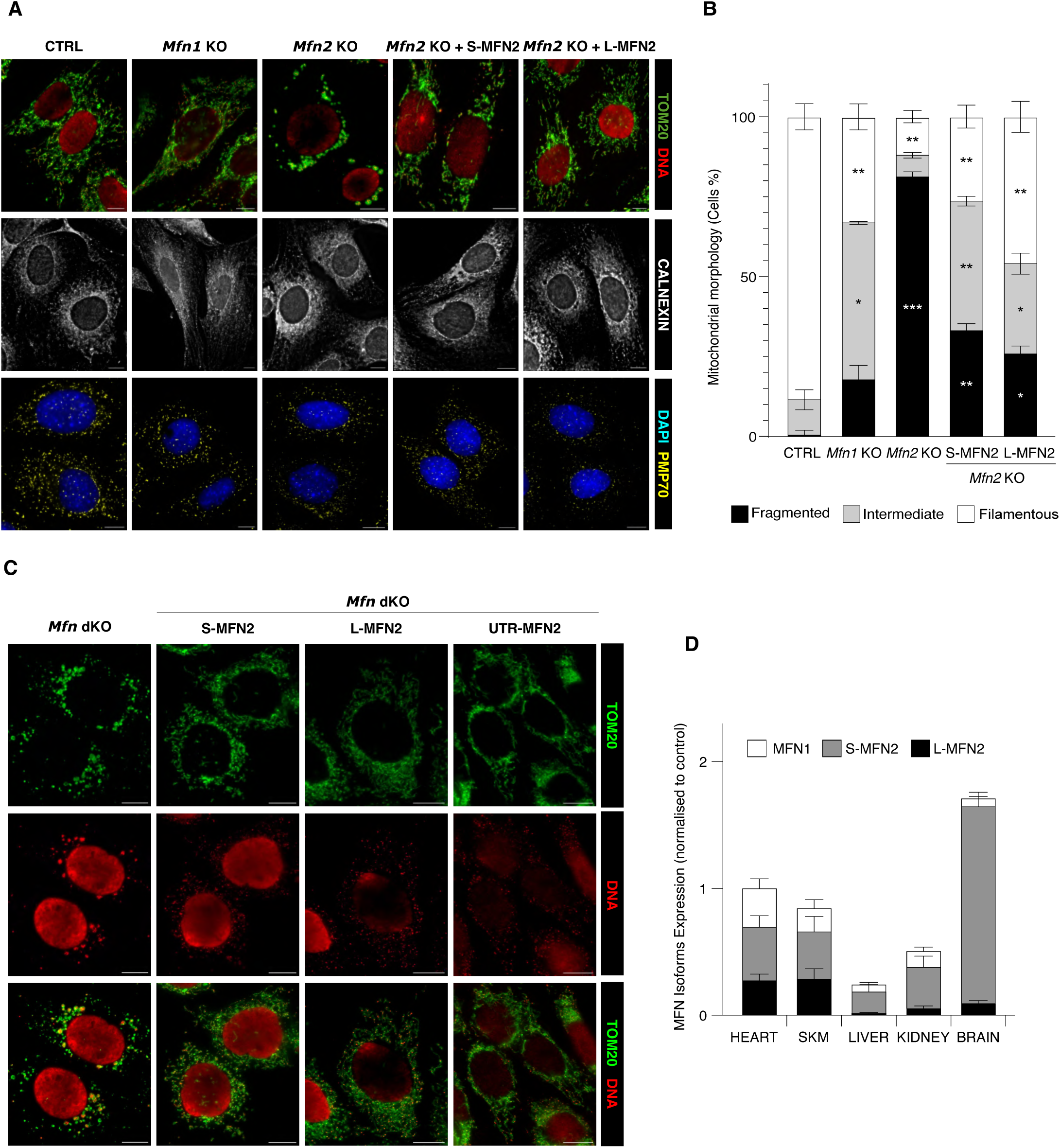
L-MFN2 and S-MFN2 mitochondrial isoforms are both fusion competent but present distinctive capacities to restore mitochondrial morphology and activity. (A) Representative epifluorescence microscopy images of the newly generated *Mfn1* KO and *Mfn2* KO MEFs expressing L-MFN2 or S-MFN2. The impact of MFN2 isoforms expression on mitochondrial (TOM20 and DNA), endoplasmic reticulum (Calnexin), and peroxisome (PMP70) morphologies was analysed using the indicated specific antibodies. (scale bar = 10 µm) (B) Quantification of mitochondrial morphology in the newly generated *Mfn* KO MEFs, expressing S-MFN2 or L-MFN2. (scale bar = 10 µm, Error bars ± SEM (*p<0.05, **p<0.005, ***p<0.0001)) (C) Representative epifluorescence microscopy of *Mfn1* and *Mfn2* dKO MEFs expressing L-MFN2, S-MFN2 or 5’UTR-MFN2. The impact of the different MFN isoforms expression on mitochondrial morphology (anti-TOM20) and mtDNA (anti-DNA) was analysed using indicated specific antibodies. (scale bar = 10 µm). (D) Densitometric quantification of the MFN1 (white), L-MFN2 (black) and S-MFN2 (grey) proportion, normalised to mitochondrial protein loaded, across the indicated mouse isolated mitochondrial extract from the indicated tissues. Quantification was achieved thank to MFN1 and MFN2 standards produced in E.coli. (n=6 mitochondrial isolations per tissue).

**Figure S6.**
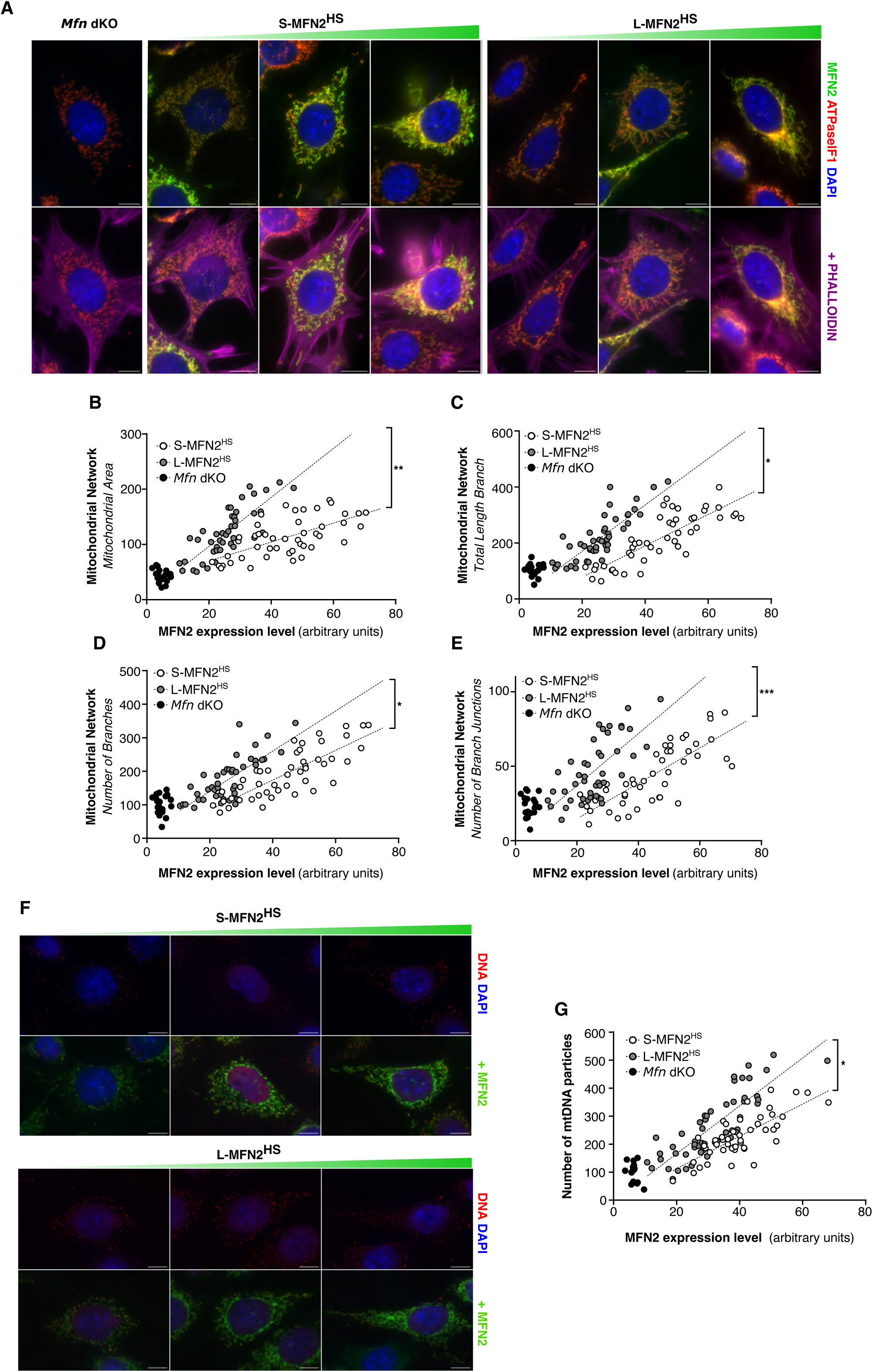
Single cell analysis of mitochondrial morphology and mitochondrial DNA particles in *Mfn* dKO MEFs lines expressing S-MFN2^HS^ or L-MFN2^HS^. (A) Representative epifluorescence microscopy of *Mfn* dKO MEFs expressing S-MFN2^HS^ or L-MFN2^HS^, using anti-ATPase IF1 as a mitochondrial marker (red), anti-MFN2 D1E9 (green), and phalloidin as a cytoskeleton marker (magenta) (scale bar = 10 µm). (B-E) Mitochondrial morphology and network parameters (mitochondrial area, total length branch, number of branches and number of branch junctions) quantified with the MitoAnalyzer plugin and plotted against the MFN2 intensity on a single-cell basis. (*n* > 30 cells for *Mfn2* KO, *Mfn2* KO + S-MFN2^HS^ and *Mfn2* KO + L-MFN2 ^HS^ MEFs respectively) PERMANOVA, the permutational multivariate analysis of variance test relative to *Mfn2* KO MEFs; *, P < 0.05; **, P < 0.01; ***, P < 0.001. (F) Representative epifluorescence microscopy of *Mfn* dKO stable MEF lines expressing S-MFN2^HS^ or L-MFN2^HS^, using anti-DNA (red) to stain mtDNA particles and anti-MFN2 D1E9 (green) (scale bar = 10 µm). (G) The number of mtDNA particles per cell is plotted against MFN2 intensity for each analyzed cell. (*n* > 30 cells for *Mfn2* KO, *Mfn2* KO + S-MFN2^HS^ and *Mfn2* KO + L-MFN2 ^HS^ MEFs respectively) PERMANOVA, the permutational multivariate analysis of variance test relative to *Mfn2* KO MEFs; *, P < 0.05; **, P < 0.01; ***, P < 0.001.

**Figure S7.**
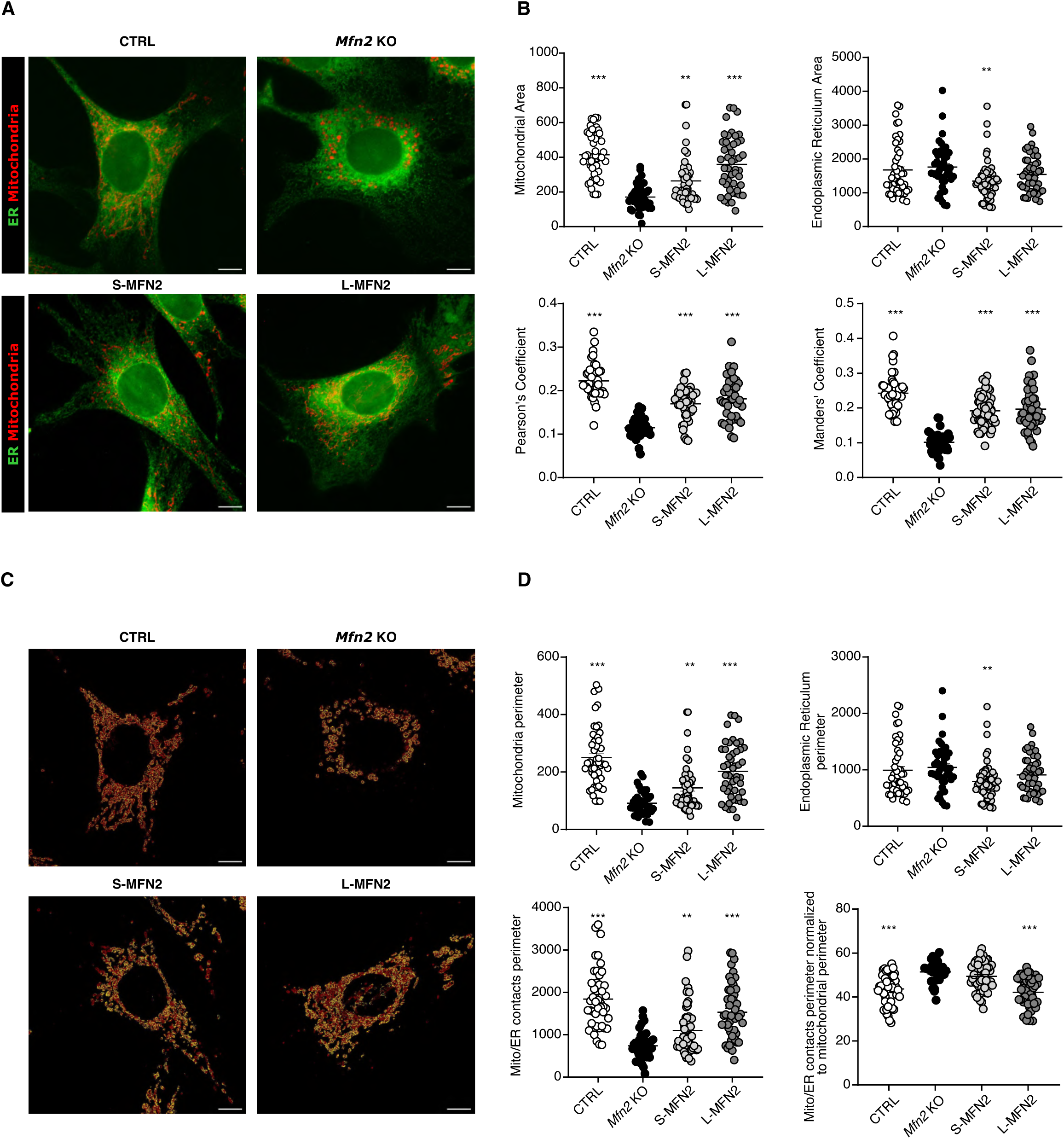
Mitochondria – Endoplasmic Reticulum Contact sites (MERCs) analysis in *Mfn2* KO MEFs expressing S-MFN2 or L-MFN2. (A) Representative epifluorescence microscopy of newly generated *Mfn2* KO MEFs expressing S-MFN2 or L-MFN2. The immunostaining was performed using the anti-calnexin as an endoplasmic reticulum marker (green) and anti-ATPase IF1 as a mitochondrial marker (red) (scale bar = 10 µm). (B) Mitochondrial area (total mitochondrial area per cell), endoplasmic reticulum area (total ER area per cell), Pearson’s, and Manders’ coefficients were scored on a single-cell basis and are represented in the plots (N = 50 *+/-5* per condition, Error bars indicate ± SEM). Statistical analyses were conducted using one-way ANOVA using Dunnett’s multiple comparison test relative to *Mfn2* KO MEFs; *, P < 0.05; **, P < 0.01; ***, P < 0.001. (C) The acquired images in A were analyzed with the IsoPhotContour2 perimeter plugin, the red pixels corresponding to the mitochondrial perimeter and the yellow pixels corresponding to the points in which the mitochondrial perimeter is in contact with the endoplasmic reticulum perimeter. (D) Mitochondrial perimeter (total mitochondrial perimeter per cell) and endoplasmic reticulum perimeter (total ER perimeter per cell) segmented using the IsoPhotContour2 perimeter plugin. The mitochondria – endoplasmic reticulum contacts perimeter per cell segmented using the IsoPhotContour2 perimeter plugin values before and after normalization to the mitochondrial perimeter are presented (N = 50 *+/-5* per condition, Error bars indicate ± SEM). Statistical analyses were conducted using one-way ANOVA using Dunnett’s multiple comparison test relative to *Mfn2* KO MEFs; *, P < 0.05; **, P < 0.01; ***, P < 0.001.

**Figure S8.**
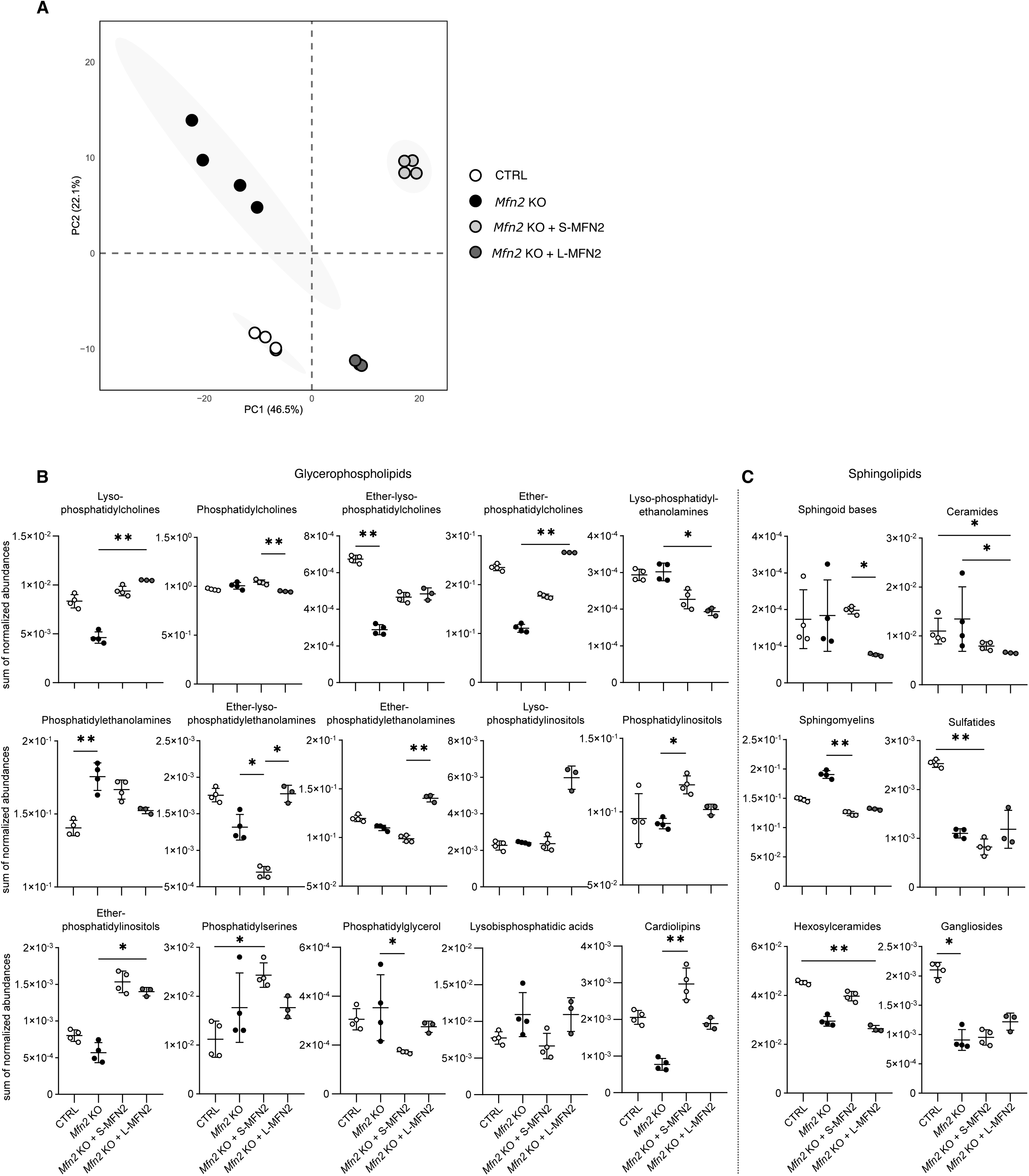
Mitochondrial lipidomic profiling in *Mfn2* KO MEFs expressing S-MFN2 or L-MFN2. (A) Scores plot from principal component analysis of mitochondrial lipidomic profiles obtained by mass spectrometry. (B) Normalized abundance of principal component analysis of glycerophospholipids profiles obtained by LC-MS (n ≥ 3, mean ± standard deviation). Statistical significance was assessed using the Kruskal-Wallis test (p<0.05) followed by post-hoc Dunn’s test (* p < 0.05, ** p < 0.01). (C) Normalized abundance of principal component analysis of Sphingolipids profiles obtained by LC-MS (n ≥ 3, mean ± standard deviation). Statistical significance was assessed using the Kruskal-Wallis test (p<0.05) followed by post-hoc Dunn’s test (* p < 0.05, ** p < 0.01).

**Figure S9.**
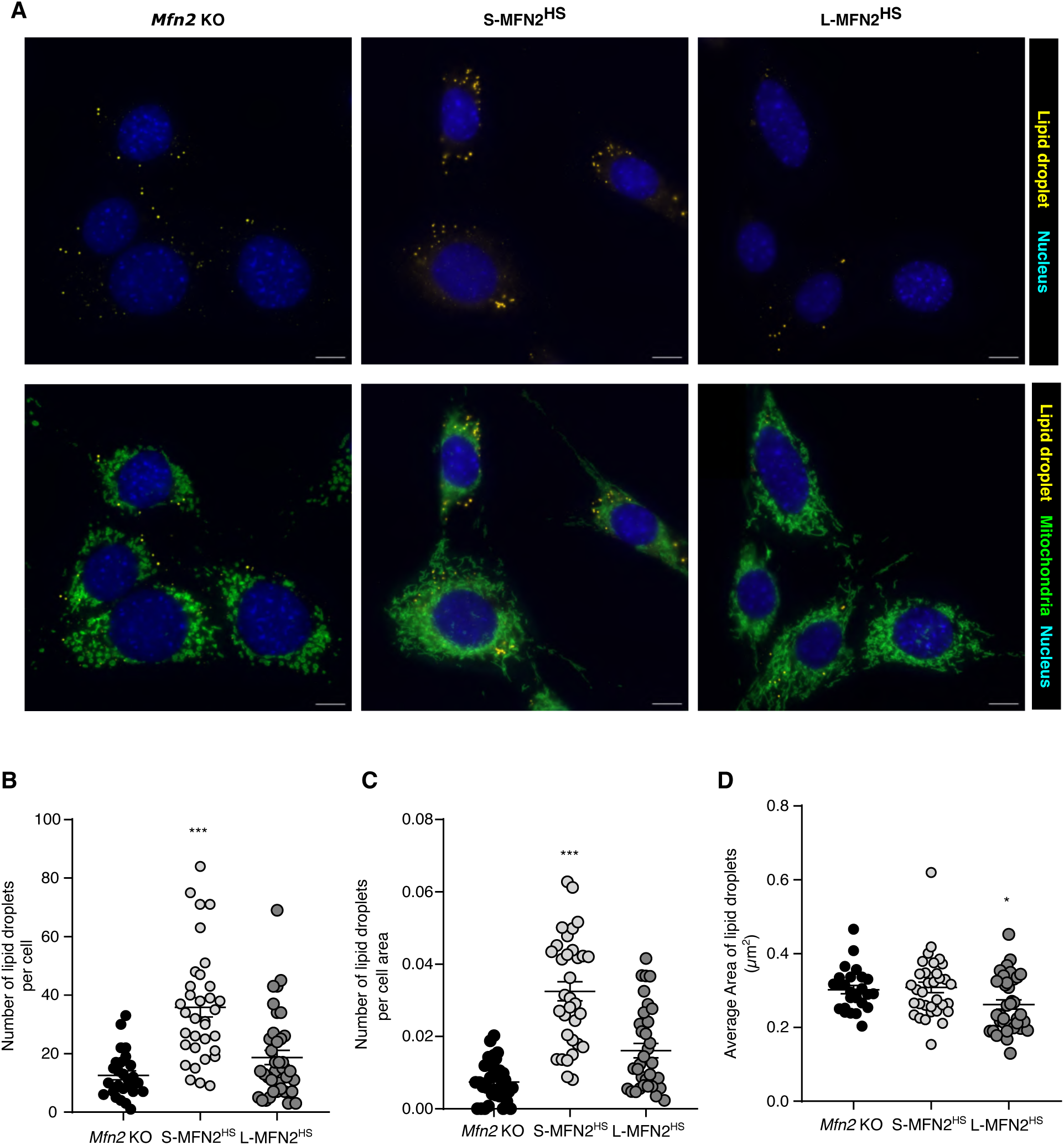
Characterization of lipid droplet levels in *Mfn2* KO MEFs expressing human S-MFN2 or L-MFN2. (A) Representative epifluorescence microscopy of *Mfn2* KO stable MEF lines expressing human S-MFN2 or L-MFN2, using NileRed as a lipid droplet marker (yellow) and anti-TOM20 as a mitochondrial marker (green) (scale bar = 10 µm). (B) The number, total (C) and average area (D) of lipid droplets were quantified on a single-cell basis in at least 50 cells. The quantification obtained are presented (*n* ≥ 30 cells for *Mfn2* KO, *Mfn2* KO + S-MFN2 and *Mfn2* KO + L-MFN2 MEFs respectively, Error bars indicate ± SEM) and subjected to statistical analysis one-way ANOVA using Dunnett’s multiple comparison test relative to *Mfn2* KO MEFs; *, P < 0.05; **, P < 0.01; ***, P < 0.001).

## Notes

### Competing Interest Statement

The authors have declared no competing interest.

